# River pollution threatens preferred macroinvertebrate prey of endemic fishes in Aotearoa, New Zealand

**DOI:** 10.64898/2025.12.08.693065

**Authors:** R.S.A. White, K.L. Hogsden, M.J. Greenwood, A.F. Rose, J. Bilewitch, P. Lambert, L. Smith, A. Brooks, O. Daly, A. Sinton, R.J Stoffels

## Abstract

1. Macroinvertebrates are a critical source of food for riverine fishes. Anthropogenic stressors can transform riverine macroinvertebrate communities, yet the consequences of such transformations to the fishes that feed on them are poorly understood. Changes in any combination of food ‘abundance,’ ‘accessibility’ or ‘quality’ may be referred to as changes in the ‘foodscape’ of fishes. Stressor-driven increases of inaccessible prey that are difficult for fish to detect, capture or ingest, may render foodscapes less profitable to fishes.
2. We examined how nutrient enrichment, fine sediment, acid mine drainage (AMD), flooding and drying reshape fish foodscapes by altering the accessibility of macroinvertebrate prey for two New Zealand river fishes, *Galaxias vulgaris* and *Gobiomorphus* breviceps. We determined relative accessibility of macroinvertebrate taxa and traits to these fishes by comparing the abundance of prey taxa and traits in the environment with their corresponding abundances in fish diets. To make these comparisons, invertebrate and fish gut samples were collected from the same river reaches, at the same times, over three years. Environmental DNA (eDNA) metabarcoding of fish gut contents facilitated efficient diet characterisation of over 700 fish guts. Finally, to determine how multiple stressors might affect fish *foodscapes*, we related invertebrate trait-specific accessibilities to published data defining how the composition of those macroinvertebrate traits varies along stressor gradients.
3. Macroinvertebrate traits that were most accessible to fishes were those promoting detection (e.g., active foraging behaviours exhibited by ‘crawlers’ and ‘scrapers’) and ingestion (e.g., moderate size, morphologies lacking defensive structures), typical of mayfly and free-living caddisfly larvae. In contrast, prey with traits inhibiting detection (e.g., small size and burrowing behaviour) and ingestion by gape-limited fishes (e.g., large size), were less accessible to the fishes we studied.
4. Trait abundance-stressor relationships revealed that that increasing fine sediment, nutrients, AMD and drying, and decreased flooding frequency, favoured macroinvertebrates with traits that erode accessibility to fishes. These results show that multiple stressors may decrease the profitability of fish foodscapes by filtering out macroinvertebrate prey with traits promoting accessibility to fishes.
5. Food web approaches to understanding multi-stressor impacts are rare, but are necessary to understand how the effects of multiple stressors on primary consumers then go on to affect predators of higher trophic position. A major barrier to fish foodscape research is the difficulty and cost of estimating trophic networks. We discuss how eDNA metabarcoding, coupled with trait-based approaches may help overcome these barriers, enabling generalisable assessments of foodscape changes caused by multiple stressors.

## Introduction

Restoration actions often focus on mitigating direct effects of stressors on fish populations (Naman et al. 2022). Far less attention is given to how environmental stressors indirectly affect fish by altering the quantity, accessibility and quality of food—the foodscape—that fish depend on (Wipfli & Baxter 2010; Naiman et al. 2012; Rossi et al. 2024). Anthropogenic stressors, such as fine sediment, nutrient enrichment and altered hydrology transform benthic macroinvertebrate communities (Dewson et al. 2007; Ardón et al. 2021; McKenzie et al. 2024), which comprise the foodscape and the basis of production for many fish species. There is, however, uncertainty about how anthropogenic changes in macroinvertebrate communities affect fish foodscapes (Ouellet et al. 2025).

A key mechanism structuring foodscapes is prey accessibility (Rossi et al. 2024; Ouellet et al. 2025). A prey species is more accessible to a predator when it is detected, captured or ingested at frequencies higher than those of other prey species. Prey accessibility is partly controlled by prey traits (Worischka et al. 2015; Rodríguez-Lozano et al. 2016). For example, nocturnal activity and reduced drift propensity can limit detection by drift-feeding fish (Rader 1997; Peckarsky & McIntosh 1998). Structural defences, including large elaborate cases, prevent ingestion (Wootton et al. 1996; Wissinger et al. 2006), while prey with large body size may exceed the mouth gape of fishes or fish life stages (Miller et al. 1988). Prey accessibility is therefore an important factor governing how shifts in prey communities affect fish growth and carrying capacity.

Understanding the controls on prey accessibility can reveal actions required to enhance fish populations. For example, environmental flow releases on the Colorado River increased trout production, despite reduced total macroinvertebrate productivity, by shifting prey from indigestible mud snails to accessible drift-active prey (Cross et al. 2011). Environmental stressors can therefore constrain or enhance fish populations by changing the relative abundance of accessible and inaccessible prey, but trophic dynamics such as these are rarely considered in environmental management.

Prey accessibility may be overlooked because it is difficult and expensive to measure (Ouellet et al. 2025) and therefore limited to relatively intensive studies in particular contexts, which in turn makes generalisations and transferability to novel contexts challenging (Green et al. 2019). A trait-based approach to understanding prey accessibility may help overcome this, noting traits are more often shared across different contexts than species (Green et al. 2019).

In this study we assessed how multiple stressors may change the foodscape of two riverine fishes in New Zealand: Canterbury galaxias (‘galaxias’; *Galaxias vulgaris*) and upland bully (‘bullies’; *Gobiomorphus breviceps*). Nutrient enrichment, fine sediment and acid mine drainage (AMD) have long degraded New Zealand rivers, typically shifting macroinvertebrate communities from those dominated by Ephemeroptera, Trichoptera, and Plecoptera (mayflies, caddisflies, and stoneflies) toward those dominated by microcrustaceans, worms, snails, or Chironomidae (Burdon et al. 2013; Graham et al. 2015; Pomeranz et al. 2019). Recent analyses show that these anthropogenic stressors, as well as flooding and drying disturbances, substantially alter the trait composition of macroinvertebrate communities (Barrett et al. 2022). We conducted an intensive study of benthic macroinvertebrate densities and fish diets over several years to estimate accessibility of prey taxa and traits to galaxias and bullies. Environmental DNA (eDNA) metabarcoding was employed to facilitate more efficient (less resource-intensive) and sensitive (more complete) characterisation of trophic networks (Elbrecht & Leese 2017). We combined results of our accessibility study with published data on how multiple stressors affect the trait composition of New Zealand’s macroinvertebrate communities to determine how stressors modify fish foodscapes. We hypothesised that nutrient enrichment, sediment, AMD and drying would increase the abundance of prey traits that are less accessible to both fish species, thereby generating a foodscape that is less profitable to bullies and galaxias.

## Methods

### Study Fish Species

Accessibility for prey taxa and traits was studied for adult and juvenile size classes of galaxias and bullies. These fishes were chosen because they are both widespread throughout the South Island of New Zealand (McDowall 1990) and often sympatric within gravel-bed rivers, but employ different feeding strategies. Canterbury galaxias are nocturnal and feed primarily on drifting macroinvertebrates (Cadwallader 1975; Glova & Sagar 1989; Glova et al. 1992), while upland bullies are diurnal benthic-feeders (Cadwallader 1975; Glova & Sagar 1989; Glova et al. 1992).

### Study Design and Sample Collection

Our study system was the Selwyn/Waikirikiri River (hereafter Selwyn), in Canterbury, New Zealand. The Selwyn River was chosen because it contains both fish species, a diverse assemblage of macroinvertebrate prey, and environmental conditions (e.g. hydrology, riparian habitat, geomorphology) representative of many rivers in New Zealand.

Two 5 ^th^ order reaches were studied: The Whitecliffs reach, which was approximately 10 km upstream of the Scotts Road reach. Each reach was 270 m long and contained nine contiguous *sampling units*, each 30 m in length. Equal amounts of macroinvertebrate and fish diet sampling was allocated to each sampling unit, to ensure our sampling was representative of the entire reach. As such, our use of sampling units helped us reduce biasing our sampling towards particular mesohabitats that, for example, might be easier to sample than others.

Sampling was completed during three *sampling years* during the austral summer (December – January): 2019-20, 2020-21, 2021-22. We wanted our estimates of prey accessibility to be representative of the summer season. To that end we sampled macroinvertebrates and fish diets each month of the summer. Within each month, three sampling units of each reach were randomly selected for sampling.

Prey accessibility was estimated by comparing macroinvertebrate prey abundance in the environment with that observed in fish diets. The abundance of prey taxa available to be consumed in the environment was estimated using benthic macroinvertebrate sampling. Within each sampling unit, thirteen benthic macroinvertebrate samples were collected per year using a Hess sampler (0.09 m^2^, 250-µm mesh). Sampling within sampling units was stratified random. First, each sampling unit was visually decomposed into sediment facies, which are units of the riverbed within which substrate composition is similar (e.g. sand; cobbles; mixed gravel/cobbles), hence habitats experiencing similar hydraulic conditions. Second, the 13 samples allocated to each sampling unit were then randomly distributed across facies. Sampling units were typically comprised of two - six facies. A total of 660 benthic samples were collected over three sampling years. All samples were preserved in 70% isopropyl alcohol on the day of collection and returned to the laboratory for processing. Taxa were identified and counted under a dissecting microscope to the lowest possible taxonomic level, which was typically species or genus, but in some cases subfamily or family (primarily Diptera or Coleoptera larvae) or class (Clitellata).

The abundance of prey taxa consumed by fish was estimated using eDNA metabarcoding of fish gut contents, which is a well developed and tested method for estimation of taxon composition in diets (Valentini et al. 2009; Pompanon et al. 2012; Deagle et al. 2019). Within each sampling unit, up to ten individuals of each fish species were captured each sampling year using backpack electrofishing. Sampling units were electrofished systematically to ensure gut samples were representative of habitats within the sampling unit. The total lengths of these fish were measured (TL, mm) before being placed immediately in 100% ethanol and returned to the laboratory for processing. Within 48 hours of collection entire digestive tracts of fish were removed. Gut contents were extracted from the digestive tract and preserved in 100% ethanol for DNA extraction and metabarcoding. At least 350 gut samples were collected for each fish species over the three sampling years.

To assess how prey accessibility varied by fish age, each fish captured was classified into either juvenile or adult age classes depending on whether its TL was greater or less than 65 mm for Canterbury galaxias, and 35 mm for upland bullies. These thresholds were based on demographic data described by Watson et al. (2025).

### eDNA Metabarcoding

The abundance of prey taxa in fish stomachs was determined through an eDNA metabarcoding approach (Pompanon et al. 2012; Deagle et al. 2019). Total DNA was extracted from each gut content sample using either a DNeasy PowerSoil Pro or a DNeasy Blood & Tissue kit (Qiagen), following the manufacturer’s protocol. DNA extracts were quantified using Quant-iT PicoGreen (Invitrogen) and 20-50ng were used in PCR targeting an approximately 300bp region of the COI mitochondrial gene using the BF1 and BR2 primers of Elbrecht and Leese (2017), with Illumina Nextera overhang adapters added to enable indexing for metabarcoding. Reactions used 1X KAPA HiFi HotStart ReadyMix and 300 nM of each primer in a 50μl total volume, with a thermocycling profile using an initial denaturation of 95 ^°^ C for 3 min, 35 cycles of 98 ^°^ C for 20 sec, 50 ^°^ C for 20 sec, and 72 ^°^ C for 10 sec, followed by a final extension of 72 ^°^ C for 1 min. PCR amplicons were visualised on 1% agarose gels using electrophoresis and successful reactions were submitted to Massey Genome Service (U. of Massey) for clean-up, indexing, library construction and 2×300bp sequencing on a MiSeq system (Illumina).

Demultiplexed sequencing reads were processed and analysed using Qiime2 v2022.2.1 (Bolyen et al. 2019). Samples were filtered, denoised, and paired reads were merged using Dada2 (Callahan et al. 2016). Taxonomic assignment was performed using two alternative reference databases constructed from GenBank (www.ncbi.nlm.nih.gov) and BoLD (Ratnasingham & Hebert 2007). The preconstructed MIDORI2 GB252 reference database (Leray et al. 2022) was used as a source of GenBank COI sequences and a customised reference database was created from BoLD COI sequences using RESCRIPt (Robeson et al. 2021). Assignment of reads against each reference database used a Naïve Bayesian classifier in Qiime2 (Bokulich et al. 2018), with each database trained on the BF1-BR2 primer region. The resulting Amplicon Sequence Variant (ASV) tables from each assignment source were examined and compared for taxonomic accuracy and resolution based on known taxa previously recovered from the studied catchments, as well as expert-identified specimens from benthic samples. Host (fish) and other vertebrate contamination was removed by filtering out reads assigned to phylum Chordata. ASVs with identical taxonomic assignments were merged and their reads were summed prior to further analysis. All bioinformatic scripts used to process the sequence data and generate reference databases are available in Appendix A1.

We defined the taxonomic resolution of a prey taxon as that at which at least ninety percent of both benthic counts and DNA reads in a particular taxonomic group (i.e., from either Class, Order, Family or Genus) could be identified to, across all samples. For example, all leptophlebiid mayfly genera were treated as separate *taxa* because over ninety percent of leptophlebiid counts were resolved to genus across both gut and benthic data. This threshold was chosen to balance detail (maximising taxonomic resolution) against reliability (avoiding misclassification due to poorly identified taxa).

### Estimation of Trait Abundance

To determine accessibility of macroinvertebrate traits we had to transform abundances of macroinvertebrate taxa into abundances of macroinvertebrate traits. Macroinvertebrate trait information was sourced from the New Zealand Freshwater Macroinvertebrate Trait Database (NZFMT) (Doledec et al. 2011). We selected five *trait groups* from the NZFMT describing the morphology and behaviour of macroinvertebrate taxa that we hypothesised would affect prey accessibility (Table 1).

**Table 1.** Macroinvertebrate prey *traits* hypothesised to influence fish predation preference. Standardised trait affinity scores for each taxon are presented in Table S1.

| *Trait group* | *Trait* | *Rationale* |
| --- | --- | --- |
| Body form (Morphology) | Cylindrical | Cylindrical and spherical taxa are often protected by cases or shells, hence less accessible. |
|  | Flattened |  |
|  | Spherical |  |
|  | Streamlined |  |
| Body flexibility (Morphology) | None (<10°) | Higher body flexibility increases handling time hence reduces accessibility. |
|  | Moderate (>10-45°) |  |
|  | High (>45°) |  |
| Maximum potential body size (Morphology) | ≤5 mm | Fish gape size limitations determine the maximum consumable prey size. |
|  | >5–10 mm |  |
|  | >10–20 mm |  |
|  | >20–40 mm |  |
|  | >40 mm |  |
| Feeding habit (Behaviour) | Algal piercer | Feeding behaviour determines prey activity and location, hence detectability. Stationary prey (e.g., filter feeders) less likely detected than active prey (e.g., predators, scrapers, shredders). |
|  | Deposit feeder |  |
|  | Filter feeder |  |
|  | Scraper |  |
|  | Shredder |  |

|  | Predator |  |
| --- | --- | --- |
| Attachment to substrate (Behaviour) | Attached | Attachment to substrate affects propensity to enter drift, or detection by benthic feeders, hence accessibility. |
|  | Burrower |  |
|  | Crawler |  |
|  | Swimmer |  |

Each *trait group* in the NZFMT is a factor containing multiple traits (Table 1). The degree to which a taxon expressed a trait was defined by *affinity scores* (Chevenet et al. (1994)). Affinity scores range [0,3], with three indicating high affinity and zero indicating no affinity, and were assigned by expert opinion (Doledec et al. 2011).

Macroinvertebrate trait abundance in a sample is simply a weighted sum, where the abundance of trait *i* in trait-group *k, t*_*ik*_, within a sample is estimated as:

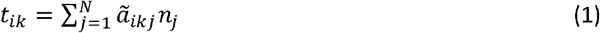

where *n*_*j*_ is the abundance of the jth taxon in the sample (there are N taxa in total), and *ã*_*ikj*_ is the standardised affinity score of the taxon for trait *i* in trait-group *k*:

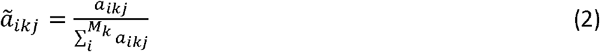

where *a*_*ikj*_ is the raw affinity score (with range [0,3]) of trait *i* for taxon *j*, and *M*_*k*_ is the number of traits in trait-group *k*. That is, standardised affinity scores are simply the raw affinity scores divided by the sum of the raw affinity scores within a trait-group (Table S1). This standardisation is required to ensure each taxon makes an equal contribution to the trait abundance estimates of a sample (Doledec et al. 2011).

### Modelling prey accessibility

Having estimated the abundance of prey taxa and traits in samples, we then modelled accessibility of each taxon and trait (hereafter, *taxon* and *trait accessibility*, respectively) to fish species and life-stages. Accessibility to a predator of taxon or trait *t* (*A*_*t*_) was quantified using Ivlev’s Electivity Index (Ivlev 1961), using the transformation of Kimmerer and Slaughter (2021). The index estimates prey accessibility by comparing the frequency of prey items eaten with their frequency in the environment:

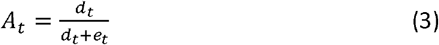

where *d*_*t*_ and *e*_*t*_ are the proportions of a taxon/trait t in the diet and the environment, respectively. *A*_*t*_ ranges [0,1], and values < 0.5 indicate consumption rates lower than expected relative to environmental availability, while values > 0.5 indicate consumption rates greater than expected relative to environmental availability.

To estimate *A*_*t*_ we had to aggregate samples. Our aggregation protocol and explanation of why we had to aggregate samples is presented in Appendix A2.

We used Mixed-Effects Hurdle Models with a beta response distribution to model the effects of stage-class on *A*_*t*_ . Our use of a model to determine *A*_*t*_ enabled us to generate more generalisable numerical estimates of A_t_ . That is, formulation of the model as *A*_*t*_ ∼ *t* + *stage-class* + *t* × stage-class + (1|*reach*/*year*) enabled us to partition the random effects of year nested within river reach from the fixed effects of interest: taxon/trait and fish stage-class (as above, t refers to either taxon or trait). We fitted this model to the data of each fish species separately.

A beta response distribution was appropriate due to the response (*trait* or *taxon accessibility*) being a continuous value between 0 and 1. However, the beta distribution cannot produce zeros, which arose when a *taxon* or *trait* was not consumed in a sample. Hurdle models are well suited to this situation because they model zeros and non-zeros by integrating two sub-models targeting different processes of data generation (Zuur et al. 2009). First, absence of a *trait* or *taxon* in the diet is modelled as a binomial process (i.e., the probability that *A*_*t*_ 0). Then, if the binomial step of the hurdle model predicts *A*_*t*_ > 0, then the beta model is used to estimate the non-zero values of *A*_*t*_. These models were fitted using the glmmTMB package in R (Brooks et al. 2017).

### Effects of environmental stressors on trait abundance

We aimed to evaluate how environmental stressors altered the abundance of traits with variable accessibilities to fishes. To do this we drew upon the analyses of Barrett et al. (2022), who analysed variation in macroinvertebrate *trait* abundance across rivers of New Zealand that spanned gradients in five stressors: flooding, drying, nutrient enrichment, fine deposited sediment, and AMD. Flooding intensity was measured using the Pfankuch River Disturbance Index (Pfankuch 1975), which reliably reflects the effects of high flow events on stream bed stability and habitability (McHugh et al. 2010; Peckarsky et al. 2014). Drying intensity was quantified as wetted cross-sectional channel area, with smaller areas reflecting greater drying stress (McHugh et al. 2015). Eutrophication was defined by a principle components analysis (PCA) axis combining nitrate and phosphorous concentrations, gross primary productivity, macrophyte cover, and shade (Graham et al. 2015), with higher scores indicating greater eutrophication stress. Fine deposited sediment was estimated visually as percent cover of particles <2 mm with higher cover considered more stressful (Burdon et al. 2013). Finally, the AMD gradient was described by a PCA axis derived from pH, conductivity, and dissolved metal concentrations (Pomeranz et al. 2019), with higher scores reflecting greater AMD stress.

Barrett et al. (2022) used multivariate ordination techniques to determine the relationships between macroinvertebrate trait abundances and levels of environmental stress. Barrett et al. (2022) presented these relationships as correlations on a continuous scale, but we used their data to present these relationships as ranked correlations between trait abundance and environmental stress, as described in detail in Appendix S3. Relationships between trait accessibility and environmental stressors were analysed graphically herein.

## Results

### Prey abundance in gut and benthic samples

A total of 703 gut contents samples were successfully amplified and DNA-sequenced, with an average of 55,165 reads produced per sample (SE=886.7, range=45-119,182). Following filtering, merging and initial processing, samples retained an average of 47,666 reads (SE=818, range=15-105,913). Taxonomic assignment (using either reference database) followed by removal of chordate ASVs resulted in an average of 30,353 reads remaining per sample (SE=934.2, range=3-96951).

The prey taxa with the highest relative abundance of DNA reads in fish guts belonged to Deleatidium, Hydropsychidae, *Hydrobiosis*, and *Coloburiscus* (Figure 1a). These taxa were disproportionately abundant in guts compared to the benthos. The *traits* with the highest relative abundance in fish diets within each *trait group* were the flattened body form, moderate body flexibility, scraper feeding habit, crawler substrate attachment and 10-20mm maximum body size (Figure 1b). The trait categories with the lowest relative abundance in fish diets within each trait were the spherical body form, no body flexibility, shredder feeding habit, swimmer substrate attachment and >40 mm maximum body size (Figure 1b).

**Figure 1:**
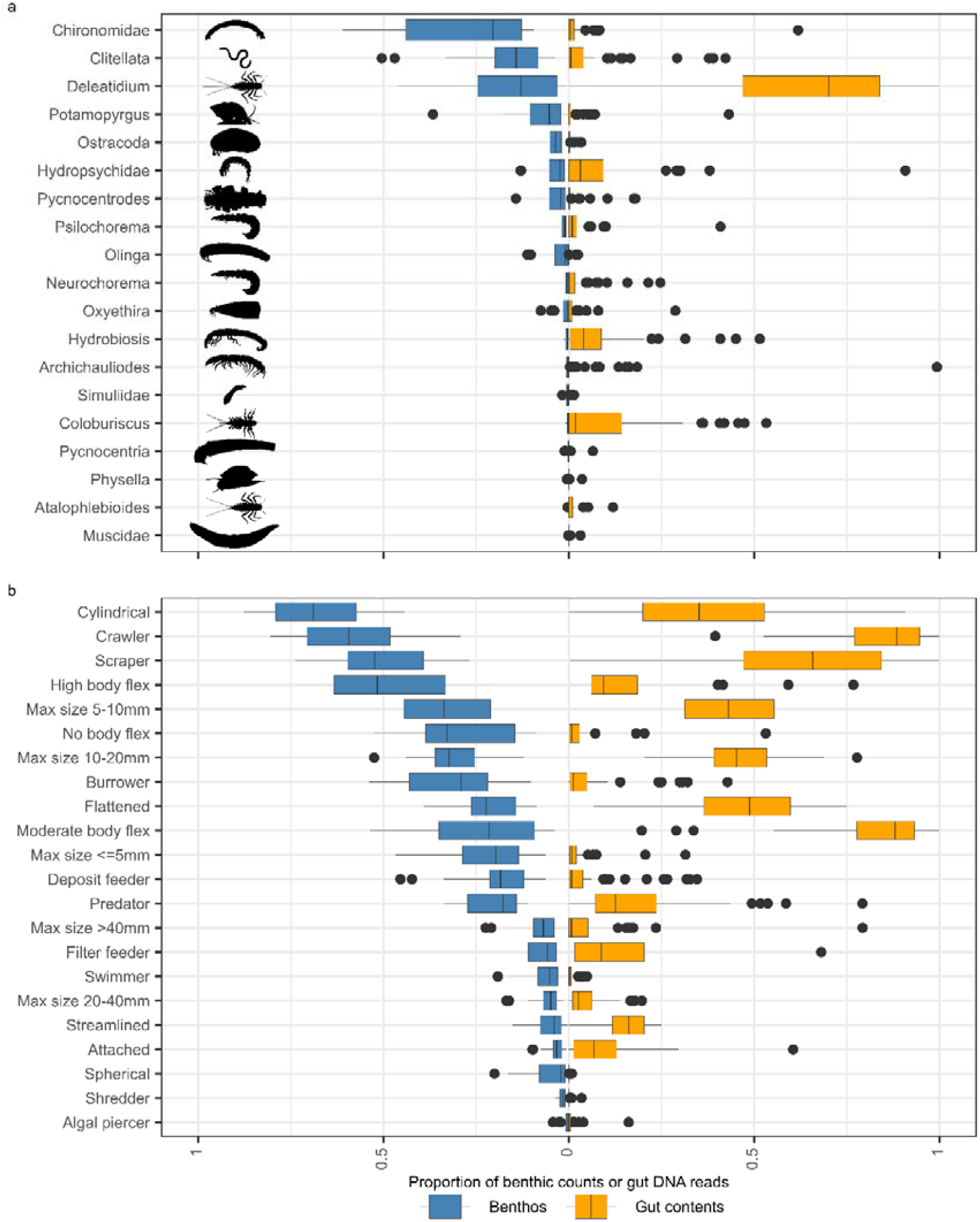
Proportion of counts in benthic samples (blue) or gut eDNA reads (orange) of a) prey taxa (y-axis) and b) prey *traits* used in models of accessibility. The box plots show the variation among all fish species, size classes, reaches and years. Silhouettes depict the approximate shape of each prey taxon made from illustrations in Winterbourn et al. (2006) where available, or sourced from https://www.phylopic.org using the *rphylopic* package (Gearty et al. 2023) (Table S2).

### Accessibility of prey taxa

*Deleatidium, Hydrobiosis*, and *Coloburiscus* were the most accessible prey taxa for adults of both fish species: *taxon accessibility* exceeded 0.5 for these prey taxa indicating they were consumed at rates greater than expected from their availability (Figure 2). Diets of juveniles of both fish species were more restricted: *taxon accessibility* exceeded 0.5 for fewer prey of juvenile fish compared to adults. Diets of juvenile bullies were the most restricted: *taxon accessibility* exceeded 0.5 for only one prey taxon – *Deleatidium* – and was less than 0.5 for all other prey taxa. Most prey taxa were only moderately or weakly accessible (*taxon accessibility* ≤ 0.5), with the lowest accessibility observed for *Olinga, Physella* and *Pycnocentria* (cased caddisflies and snails) (Figure 2).

**Figure 2:**
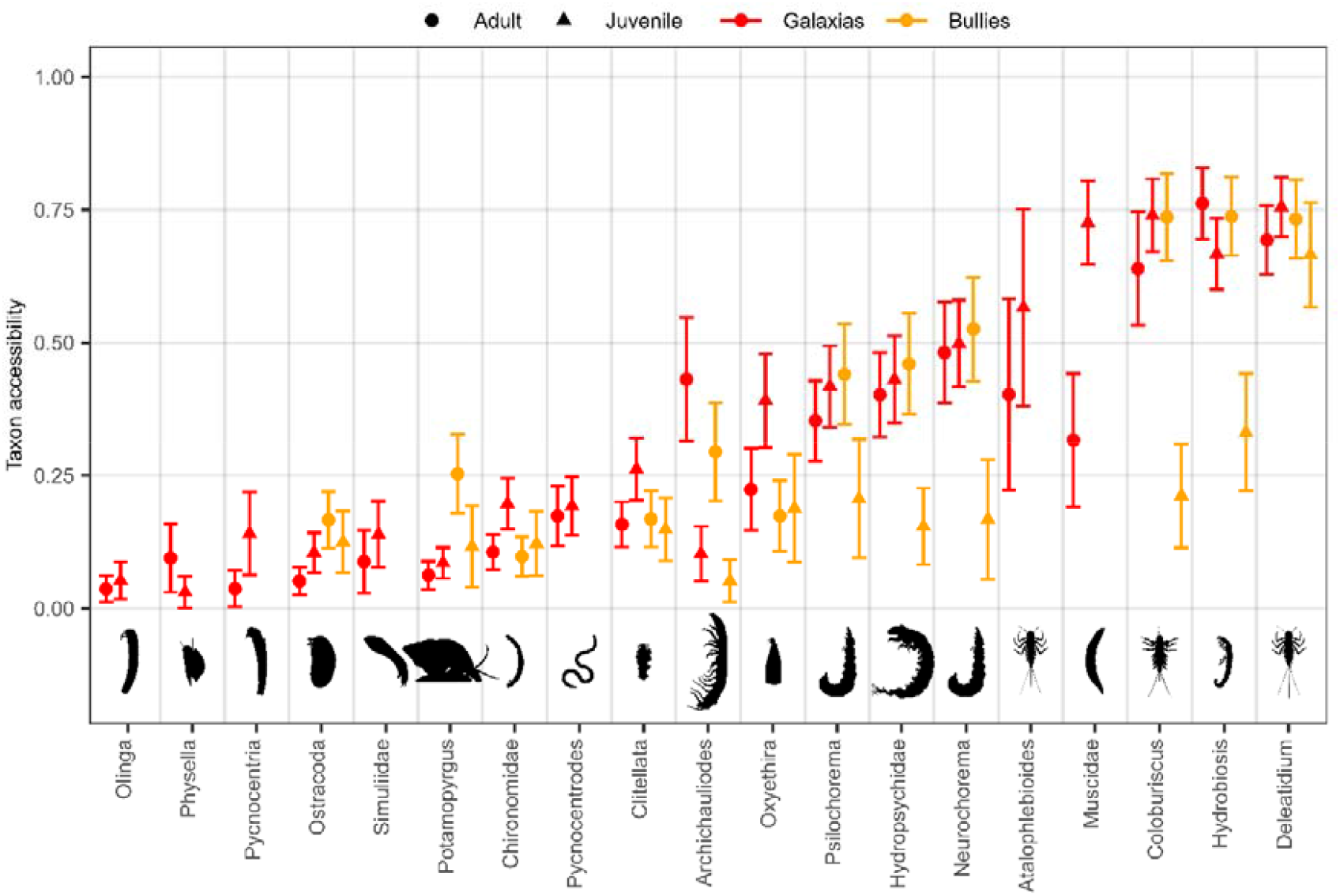
*Taxon accessibility* for adult (circles) and juvenile (triangles) Canterbury galaxias (red), and upland bully (orange) as predicted by the hurdle models of *taxon accessibility* for each fish species (mean points, 95% confidence interval error bars). Taxa that were consumed in less than two sample occasions for a particular predator and age class are not shown.

### Accessibility of prey traits

*Trait accessibility* for all fishes was highest for traits typically characterised by mayflies and free-living caddisflies: moderate body flexibility, flattened or streamlined body forms, crawler substraten attachment and scraper feeding habit (Figure 3a). Accessibility for all fish species and size classes was highest for moderate size prey (10-20 mm) (Figure 3c). The least accessible traits for all fishes included size extremes (≤5mm and >40mm), body flexibility extremes (no and high body flexibility), spherical and cylindrical body forms, shredder and deposit feeders, and swimmer substrate attachment (Figure 3a). These traits were most characteristic of microcrustaceans, cased-caddisflies, snails, and clitellates, which were among the least accessible taxa (Figure 2). *Trait accessibility* for juveniles of both fish species more restricted compared to adults, particularly for prey with cylindrical body shape, and predator and filter-feeder feeding habits, attached substrate attachment (Figure 2). Accessibility of large prey to juvenile upland bullies (>20 mm) was notably lower than that of adults (Figure 3c).

**Figure 3:**
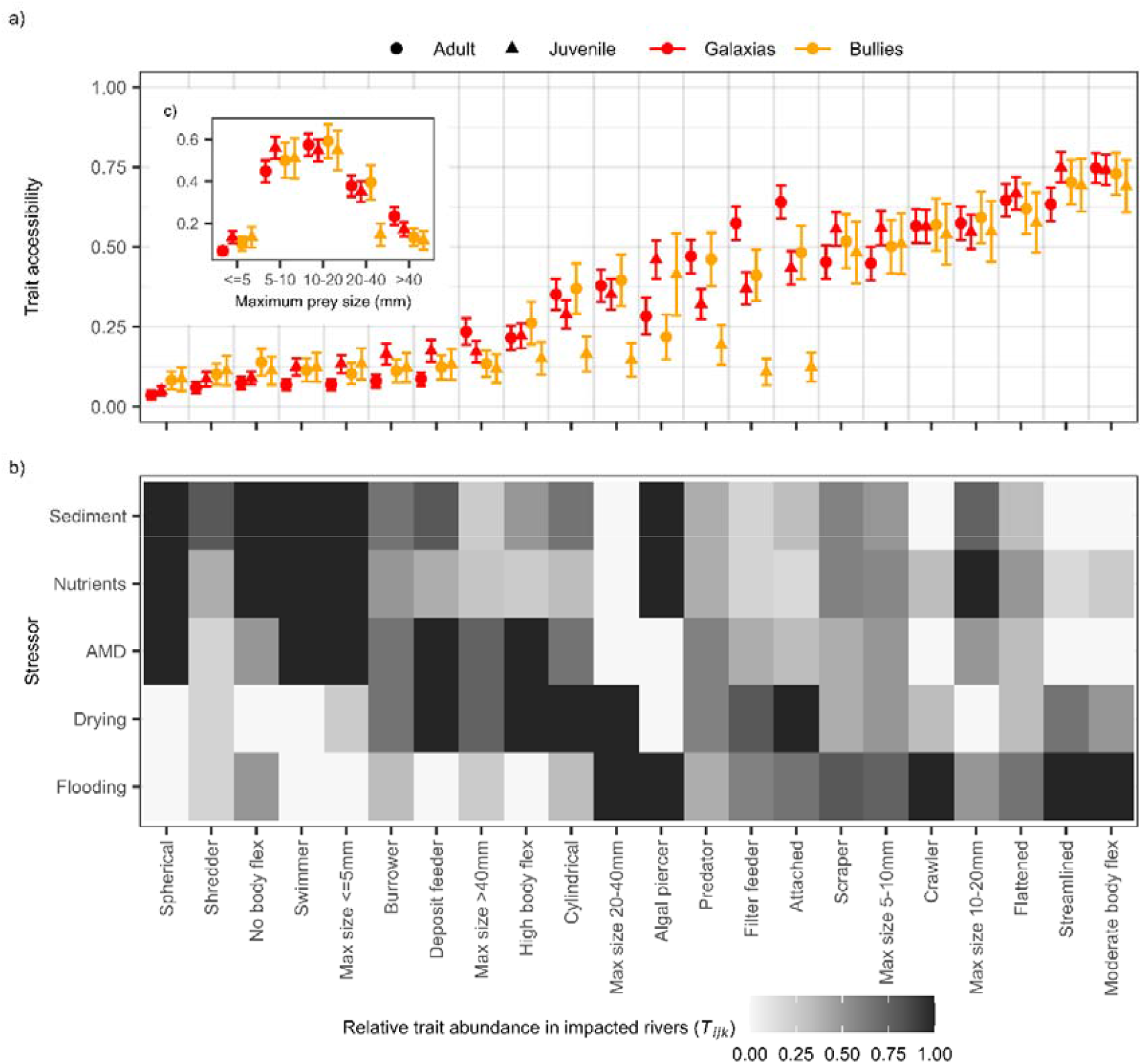
The association between a) *trait accessibility* and b) relative trait abundance in rivers impacted by stressors (*T*_*iks*_). Inset panel c) shows the relationship between *trait accessibility* and prey size. *Trait accessibility* estimates from hurdle models for adult (circles) and juvenile (triangles) Canterbury galaxias (red) and upland bully fish (mean points, 95% confidence interval error bars). *Traits* in a) and b) are ordered left to right from the least to most accessible on average among both fishes and age classes. High *T*_*iks*_ values (greyscale) indicate the trait is highly abundant under stressful conditions of a stressor (i.e., high deposited fine sediment cover, nutrient enrichment, AMD and larger and more frequent drying and flooding events) relative to other *traits* within a *trait group*.

### Distribution of accessible prey traits among environmental stressors

The flooding stressor was associated with dominance of macroinvertebrate *traits* that were highly accessible to both fish predators: moderate flexibility, streamlined body forms and crawlers characteristic of mayflies, as well as moderately accessible large sized (20-40 mm) and algal piercing prey (Figure 3b). The drying stressor was generally associated with a reduction in dominance of the most accessible traits and increased dominance of moderately accessible traits. The deposited fine sediment, nutrient enrichment and AMD stressors were more associated with dominance of prey *traits* that were least accessible as prey by both fishes: small body size (≤ 5 mm), spherical shape, swimmers, and no body flexibility, characteristic of microcrustaceans and snails). Traits associated with clitellate worms (deposit feeder, high flexibility) were also had high dominance in AMD and sediment impacted streams.

## Discussion

Canterbury galaxias and upland bully accessed a narrow range of prey traits relative to those available in the environment. Macroinvertebrate traits associated with high accessibility scores were morphological traits that may facilitate ingestion by the relatively small invertivorous fishes studied here—moderate size (>5 mm and < 20 mm) and bodies without defensive cases and shells (e.g. streamlined, flattened morphologies)—and behavioural traits that may facilitate prey detection by fishes (crawlers, scrapers, predators and filter feeders). These traits were characteristic of mayfly larvae—which comprised over 70% of gut contents despite representing only 13% of prey in the environment—and uncased or ‘free-living’ caddis larvae.

In contrast, macroinvertebrate traits less accessible to fishes were morphological traits that arguably inhibit ingestion–large size (> 20 mm) and protective shells or cases (e.g. spherical shape, rigid bodies)—and behavioural and morphological traits that reduce detection—small size (< 5 mm) and a tendency to inhabit small interstitial spaces of the benthos (burrowers and deposit-feeders). These traits were characteristic of snails, worms and chironomids.

When we linked these accessibility results with trait abundance-stressor relationships, we found that increasing fine sediment, nutrient concentrations and AMD tend to favour the macroinvertebrate traits less preferred or less accessible to galaxias and bullies. With respect to hydrological variables, increased river drying and decreased flooding also tended to favour macroinvertebrate traits that are less accessible to the invertivorous fishes of our study. Thus, multiple anthropogenic stressors may transform fish foodscapes to ones comprised of less accessible prey, hence foodscapes potentially less profitable to some invertivorous fishes.

### Traits affecting prey accessibility

The patterns of *trait accessibility* we observed reflected mechanisms controlling prey detection, capture and ingestion (Ouellet et al. 2025). For benthic and drift-feeding fishes—like galaxias and bullies—prey traits associated with higher drift propensity and visibility likely increase detectability and capture. For example, scrapers were the most accessible feeding groups probably because they feed on algae on top of rocks, where they are visible to benthic predation and exposed to shear stresses that may dislodge individuals and cause involuntary drifting (Rader 1997; De Crespin De Billy & Usseglio-Polatera 2002). In contrast, some deposit-feeders and shredders have burrowing habits, and so may remain hidden within sediments or coarse organic matter (Rader 1997; Herbst et al. 2018), where they are less accessible to fishes (Suttle et al. 2004).

Once detected, traits that reinforce attachment to substrates may reduce ingestion efficiency. Such traits include use of silk threads and suction cups by dipterans (Fingerut et al. 2006; Kang et al. 2021). This may explain why attached prey like Simuliidae were less accessible to our fishes than macroinvertebrates with, for example, crawling attachment—crawlers may be more easily dislodged, hence ingested (Rader 1997).

Prey size also mediates detection, capture and ingestion. Larger prey are more easily detected, but harder to capture and ingest due to gape size limitations of fishes (Werner & Hall 1974; Persson & Brönmark 2002). Smaller prey, however, provide less energetic returns and may be overlooked if larger prey are detected (Werner & Hall 1974). These mechanisms can result in hump-shaped relationships between prey accessibility and size—intermediate sized prey present an optimal balance between detectability, ease of capture, ingestion, and energetic return (Werner & Hall 1974; Wahlström et al. 2000; Persson & Brönmark 2002). Accordingly, we found intermediate sized (10-20mm) prey were the most accessible, while size extremes (≤5mm, >40mm) were the least accessible. Optimal prey sizes also depended on predator size, particularly for bullies: accessibility of prey >20 mm was significantly lower for juvenile bullies compared to adults. This likely reflects the stronger constraints of gape size in juvenile fish (Miller et al. 1988).

Prey accessibility patterns generally aligned with established trade-offs between traits that promote tolerance of predation and hydraulic stress (Wootton et al. 1996). Macroinvertebrate traits of low accessibility—small, inflexible, cylindrical-shaped morphologies – were characteristic of snails (e.g., *Potamopyrgus*) and cased-caddisflies (e.g., *Olinga*). Large cases of such prey often defend against ingestion (Wissinger et al. 2006; Limm & Power 2011), but also increase hydraulic drag, restricting them to lower velocities (Wootton et al. 1996; Verhaegen et al. 2019). In contrast, the more accessible prey—mayflies and free-living caddisflies—are more streamlined or flattened and tolerant of higher velocities (Collier 1994; Townsend et al. 1997). These prey may be easier to detect and capture due to their moderate size, limited defences, and higher drift-propensity (Rader 1997).

Many prey traits included in our analysis may have been somewhat redundant because they correlated with other traits that shared similar mechanisms driving accessibility. For example, deposit-feeders and burrowers, were the least accessible substrate-attachment and feeding-habit traits, probably because they are both behaviours that hide prey from predators (Rader 1997). Redundancy may also arise due to spurious correlations with other traits that are the more likely driving mechanism affecting accessibility. For example, inflexible-bodied prey were among the least accessible, even though such prey might be less capable of evading capture (Domenici et al. 2011). However, low body flexibility was also associated with prey having rigid shells or cases, which often protect prey against ingestion (Wootton et al. 1996). Consequently, the low accessibility of inflexible taxa was likely driven by these defensive structures, rather than flexibility of the bodies inside those structures.

### Environmental stressors restructure foodscapes via trait filtering

Anthropogenic pressures have been mounting in the world’s rivers for decades (Dudgeon et al. 2006; Vörösmarty et al. 2010; Reid et al. 2019). Flow regimes of about two-thirds of Earth’s large rivers have been altered by dams and reservoirs (Grill et al. 2019); nitrogen concentrations have increased by 35% on average, doubling the number of rivers impacted by harmful algal blooms (McDowell et al. 2025); while erosion from land-use change has increased fine sediment pollution in many catchments (Dethier et al. 2022). The impacts of these stressors on macroinvertebrate communities are relatively well known (Dewson et al. 2007; Ardón et al. 2021; McKenzie et al. 2024), with losses of sensitive taxa reshaping communities (Stoffels & White 2024). The extensive evidence showing how multiple stressors reshape macroinvertebrate communities is compelling (Birk et al. 2020), but begs the question: what are the consequences for food web processes and, in particular, fish foodscapes? Our study makes some progress towards answering this question, and shows that multiple anthropogenic stressors may erode the profitability of fish foodscapes by filtering out traits promoting accessibility to fishes.

Our findings reveal mechanisms by which stressors reshape fish foodscapes through changes in macroinvertebrate communities. For example, the dominance of inaccessible prey in sediment-impacted rivers likely reflects habitat-size constraints from infilling of interstitial spaces (Burdon et al. 2013). These constraints exclude larger accessible prey, such as *Deleatidium* (Burdon et al. 2013), while favouring burrowing deposit feeders that are physically unavailable to fish, and lead to reduced fish growth and recruitment (Suttle et al. 2004). Similarly, thick metal hydroxide deposits in AMD streams favour deposit-feeding worms (oligochaetes) and chironomids, while excluding accessible grazers such as *Deleatidium* (Hogsden & Harding 2012; Hogsden et al. 2013), contributing to fish density declines downstream of mines (Scullion & Edwards 1980). Meanwhile, in nutrient-enriched streams, reduced foodscape quality primarily resulted from proliferation of predation-resistant grazers—the New Zealand mudsnail—which proliferate on macrophyte surfaces (Graham et al. 2015). The increased dominance of accessible, streamlined prey in flood-disturbed rivers likely reflects flow-related filters selecting for velocity tolerance (Wootton et al. 1996). This helps explain why fish foodscapes may shift in regulated rivers where flood frequencies or magnitudes have been reduced: selective pressures that exclude less profitable prey are reduced (Wootton et al. 1996; Cross et al. 2011).

### Potential strengths and limitations of our approach

If foodscape approaches are to more commonly inform freshwater ecosystem management then we require cost-effective, efficient ways to characterise trophic networks (Naman et al. 2022; Ouellet et al. 2025). Estimating the diet composition of predators can be a laborious and challenging process, historically involving the examination of partially digested prey items extracted from predator guts. In the present study we employed eDNA metabarcoding to estimate the taxon composition of several hundred diets. This technique saves labour costs (McInerney & Rees 2018), and may decrease misidentification of consumed taxa which, having undergone partial mechanical and chemical digestion, can be hard to identify based on morphological features (Valentini et al. 2009; Pompanon et al. 2012; Joly et al. 2014). The potential of eDNA technologies to improve the efficiency and effectiveness of ecological network research has recently been noted (Evans et al. 2016), but at this stage few freshwater scientists have applied these technologies to applied foodscape studies. Molecular biologists are rapidly developing eDNA technologies, such that we may move beyond detection of species’ presence-absence in samples, to estimation of species relative abundance and biomass (Elbrecht & Leese 2015; Thomas et al. 2016; Bista et al. 2018; Schenk et al. 2019). These technologies may play a key role in normalising foodscape approaches to water resource management.

Fishes often exhibit frequency-dependent predation, such that their prey preferences vary in space and time to correlate positively with whichever prey taxa are most abundant (Falke et al. 2020). Experiments have shown that when alternative prey with equal accessibilities vary in abundance, fishes switch their preferences to prey that are most abundant (Murdoch et al. 1975; Hughes & Croy 1993). What is less clear is the extent to which frequency-dependent predation is present in natural, complex systems where prey do not just vary in abundance, but also in accessibility, where accessibilities are driven by traits of prey, as well as foraging traits of the predator (Allen et al. 1988; Prokopenko et al. 2023). Given our modelling approach we have effectively assumed that prey accessibilities to fish are a relatively fixed property of the ecology of fish-macroinvertebrate interactions, mediated by traits that fundamentally limit accessibility. A useful extension to our work would be an investigation of predatory-prey systems across numerous spatial and temporal contexts, to determine how sensitive prey accessibilities are to context (Greenwood 2008). Such studies will be key to generalising and validating the inferences we present herein.

## Conclusion

We have shown that multiple anthropogenic stressors may erode the profitability of fish foodscapes by filtering out macroinvertebrates with traits that enhance their accessibility to some fishes. Studies such as this play a critical role in improving our understanding of how transformation of freshwater macroinvertebrate communities affects vertebrate consumers higher in the food web. Foodscape approaches to freshwater ecosystem assessment may become more routine as freshwater ecologists collaborate with molecular ecologists to integrate eDNA approaches into food web studies.

## Acknowledgements

This work was funded through the Ministry of Business, Innovation and Employment’s Strategic Science Investment Fund. We gratefully acknowledge Michael Winterbourn for permission to use his illustrations and Simon Hayes for tracing the macroinvertebrate silhouettes from them.

## Data availability statement

Data are available from the authors upon reasonable request.

### Author contribution statement

Conceptualisation: RSAW, RJS, KH. Developing methods: RSAW, RJS, KH, AR, JB, PL, LS, AB, OD, AS. Conducting the research: RSAW, RJS, KH, MG, AR, JB, PL, LS, AB, OD, AS. Data analysis: RSAW, RJS. Data interpretation: RSAW, RJS, MG, KH. Preparation figures and tables: RSAW, RJS, KH. Writing: RSAW, RJS, MG, KH.

