## Appendices for "River pollution threatens preferred macroinvertebrate prey of endemic fishes in Aotearoa, New Zealand"

**Supplementary Information**

**Appendix A1**

a) Qiime2 script for processing MiSeq metabarcoding sequence data from fish gut contents

qiime tools import \

--type 'SampleData[PairedEndSequencesWithQuality]' \

--input-path fastq \

--input-format CasavaOneEightSingleLanePerSampleDirFmt \

--output-path 1_FastQ_Imported.qza

qiime dada2 denoise-paired \

--i-demultiplexed-seqs 1_FastQ_Imported.qza \

--p-trim-left-f 50 \

--p-trim-left-r 50 \

--p-trunc-len-f 220 \

--p-trunc-len-r 160 \

--p-min-overlap 12 \

--p-chimera-method consensus \

--o-table 2_Dada2_Table.qza \

--o-representative-sequences 2_Dada2_RepSeqs.qza \

--o-denoising-stats 2_Dada2_DenoiseStats.qza

qiime feature-classifier classify-sklearn \

--i-classifier ~/RefDb/refdb_BOLD_COI/bold_classifier_BF1BR2.qza \

--i-reads 2_Dada2_RepSeqs.qza \

--o-classification 3_Taxonomy_BOLD.qza

qiime feature-classifier classify-sklearn \

--i-classifier ~/RefDb/refdb_Midori_COI_NEW/midori_derep_classifier.qza \

--i-reads 2_Dada2_RepSeqs.qza \

--o-classification 3_Taxonomy_Midori.qza

qiime taxa filter-table \

--i-table 2_Dada2_Table.qza \

--i-taxonomy 3_Taxonomy_BOLD.qza \

--p-exclude p__Chordata \

--o-filtered-table 4_Table-noChord_BOLD.qza

qiime taxa filter-table \

--i-table 2_Dada2_Table.qza \

--i-taxonomy 3_Taxonomy_Midori.qza \

--p-exclude p__Chordata \

--o-filtered-table 4_Table-noChord_Midori.qza

qiime taxa collapse \

--i-table 4_Table-noChord_BOLD.qza \

--i-taxonomy 3_Taxonomy_BOLD.qza \

--p-level 7 \

--o-collapsed-table 4_Table-noChord_BOLD.qza

qiime taxa collapse \

--i-table 4_Table-noChord_Midori.qza \

--i-taxonomy 3_Taxonomy_Midori.qza \

--p-level 7 \

--o-collapsed-table 4_Table-noChord_Midori.qza

b) Customised COI reference database creation from BoLD

All initial ‘R’ code for BoLD data downloading and initial pruning plus assembly was obtained from <https://forum.qiime2.org/t/building-a-coi-database-from-bold-references/16129> (Robeson et al. 2021). Remainder of code was adapted from the same source, for implementation in Qiime2:

qiime tools import \

--input-path 3_bold_rawSeqs_forQiime_noGaps.fasta \

--output-path 3_bold_rawSeqs.qza \

--type 'FeatureData[Sequence]'

qiime tools import \

--type 'FeatureData[Taxonomy]' \

--input-format HeaderlessTSVTaxonomyFormat \

--input-path 3_bold_rawTaxa_forQiime.tsv \

--output-path 3_bold_rawTaxa.qza

qiime rescript cull-seqs \

--i-sequences 3_bold_rawSeqs.qza \

--p-num-degenerates 5 \

--p-homopolymer-length 12 \

--o-clean-sequences 4_bold_rawSeqs_culled.qza

qiime rescript dereplicate \

--i-sequences 4_bold_rawSeqs_culled.qza \

--i-taxa 3_bold_rawTaxa.qza \

--p-mode 'super' \

--p-derep-prefix \

--p-rank-handles 'greengenes' \

--o-dereplicated-sequences 4_bold_derep_seqs.qza \

--o-dereplicated-taxa 4_bold_derep_taxa.qza

\# filter taxonomy, then seqs for anything not IDd to at least phylum

qiime rescript filter-taxa \

--i-taxonomy 4_bold_derep_taxa.qza \

--p-exclude 'p__;' \

--o-filtered-taxonomy 4_bold_derep_filt_taxa.qza

qiime taxa filter-seqs \

--i-sequences 4_bold_derep_seqs.qza \

--i-taxonomy 4_bold_derep_taxa.qza \

--p-exclude 'p__;' \

--o-filtered-sequences 4_bold_derep_filt_seqs.qza

qiime feature-classifier extract-reads \

--i-sequences 4_bold_derep_filt_seqs.qza \

--p-f-primer ACWGGWTGRACWGTNTAYCC \

--p-r-primer TCDGGRTGNCCRAARAAYCA \

--p-n-jobs 10 \

--o-reads 5_bold_ref-seqs_BF1BR2.qza

qiime rescript dereplicate \

--i-sequences 5_bold_ref-seqs_BF1BR2.qza \

--i-taxa 4_bold_derep_filt_taxa.qza \

--p-mode 'uniq' \

--p-threads 20 \

--o-dereplicated-sequences 5_bold_ref-seqs_BF1BR2_derep.qza \

--o-dereplicated-taxa 5_bold_derep_filt_taxa_derep.qza

qiime rescript cull-seqs \

--i-sequences 5_bold_ref-seqs_BF1BR2_derep.qza \

--p-n-jobs 10 \

--p-num-degenerates 1 \

--p-homopolymer-length 8 \

--o-clean-sequences 5_bold_ref-seqs_BF1BR2_derep_cull.qza

qiime rescript extract-seq-segments \

--i-input-sequences 4_bold_derep_filt_seqs.qza \

--i-reference-segment-sequences 5_bold_ref-seqs_BF1BR2_derep_cull.qza \

--p-perc-identity 0.7 \

--p-min-seq-len 10 \

--o-extracted-sequence-segments 6_bold_extracted_refs-01.qza \

--o-unmatched-sequences 6_bold_extracted_unmatchedseqs-01.qza

qiime rescript dereplicate \

--i-sequences 6_bold_extracted_refs-01.qza \

--i-taxa 4_bold_derep_filt_taxa.qza \

--p-mode 'uniq' \

--o-dereplicated-sequences 6_bold_extracted_refs_derep-01.qza \

--o-dereplicated-taxa 6_bold_derep_filt_taxa_derep-01.qza

qiime rescript cull-seqs \

--i-sequences 6_bold_extracted_refs_derep-01.qza \

--p-num-degenerates 1 \

--p-homopolymer-length 8 \

--o-clean-sequences 6_bold_extracted_refs_derep_cull-01.qza

\## 2nd iteration to include refseq capture

qiime rescript extract-seq-segments \

--i-input-sequences 4_bold_derep_filt_seqs.qza \

--i-reference-segment-sequences 6_bold_extracted_refs_derep_cull-01.qza \

--p-perc-identity 0.7 \

--p-min-seq-len 10 \

--o-extracted-sequence-segments 7_bold_extracted_refs-02.qza \

--o-unmatched-sequences 7_bold_extracted_unmatchedseqs-02.qza \

--verbose

qiime rescript dereplicate \

--i-sequences 7_bold_extracted_refs-02.qza \

--i-taxa 4_bold_derep_filt_taxa.qza \

--p-mode 'uniq' \

--o-dereplicated-sequences 7_bold_extracted_refs_derep-02.qza \

--o-dereplicated-taxa 7_bold_derep_filt_taxa_derep-02.qza

qiime rescript cull-seqs \

--i-sequences 7_bold_extracted_refs_derep-02.qza \

--p-num-degenerates 1 \

--p-homopolymer-length 8 \

--o-clean-sequences 7_bold_extracted_refs_derep_cull-02.qza

\## final trimming

qiime rescript filter-seqs-length \

--i-sequences 7_bold_extracted_refs_derep_cull-02.qza \

--p-global-min 10 \

--p-global-max 1000 \

--o-filtered-seqs 7_bold_extracted_refs_derep_cull_keep-02.qza \

--o-discarded-seqs 7_bold_extracted_refs_derep_cull_discard-02.qza

qiime rescript filter-taxa \

--i-taxonomy 4_bold_derep_filt_taxa.qza \

--m-ids-to-keep-file 7_bold_extracted_refs_derep_cull_keep-02.qza \

--o-filtered-taxonomy 7_bold_extracted_taxa_derep_cull_keep-02.qza

qiime feature-classifier fit-classifier-naive-bayes \

--i-reference-reads 7_bold_extracted_refs_derep_cull_keep-02.qza \

--i-reference-taxonomy 7_bold_extracted_taxa_derep_cull_keep-02.qza \

--o-classifier bold_classifier_BF1BR2.qza

### c) Read counts per sample at each stage of DNA sequence processing.

‘Raw’ = # demultiplexed reads prior to filtering, ‘Filtered’ = # reads post-filtering, ‘Denoised’ = # reads post-denoising (via Dada2), ‘Merged’ = # R1 and R2 (paired-end) reads joined, ‘Non-chimeric’ = # joined reads remaining after removal of putative chimeric sequences, ‘Final ASVs’ = # taxonomy-assigned reads (ASVs) remaining after removal of chordate sequences. Sample replicates are denoted by a ‘.1’ and ‘.2’ suffix. Suffixes for negative controls (‘NEG’) indicate sequencing run (1, 2, 3) followed by a replicate number (for NEG_3 only).

| **Sample #** | **Raw** | **Filtered** | **Denoised** | **Merged** | **Non-chimeric** | **Final ASVs** |
| --- | --- | --- | --- | --- | --- | --- |
| 1 | 42295 | 36193 | 35617 | 35484 | 35474 | 1627 |
| 2 | 45949 | 39583 | 38872 | 38685 | 36543 | 1401 |
| 3 | 40936 | 35422 | 35100 | 35035 | 34940 | 27171 |
| 4 | 43125 | 36931 | 36414 | 36014 | 33764 | 22306 |
| 5 | 52293 | 40502 | 39495 | 39026 | 39022 | 1995 |
| 6 | 43543 | 38151 | 37586 | 37422 | 37327 | 22015 |
| 7 | 39940 | 35192 | 34847 | 34673 | 34022 | 22001 |
| 9 | 44035 | 37860 | 36975 | 36584 | 35545 | 600 |
| 10 | 44335 | 38993 | 38694 | 38525 | 38171 | 21020 |
| 11 | 47852 | 41235 | 40687 | 40177 | 39498 | 39431 |
| 12 | 50777 | 43926 | 42821 | 41899 | 41821 | 34405 |
| 13 | 41768 | 35077 | 34084 | 33292 | 32932 | 30796 |
| 14 | 48166 | 42390 | 41957 | 41761 | 41704 | 41556 |
| 17 | 47325 | 41828 | 41050 | 40319 | 39573 | 39279 |
| 18 | 52797 | 45372 | 44922 | 44659 | 37463 | 37323 |
| 19 | 45005 | 39302 | 38275 | 37313 | 34937 | 31786 |
| 20 | 45409 | 38317 | 37432 | 36766 | 33523 | 11381 |
| 21 | 45419 | 39830 | 39346 | 39207 | 39075 | 18330 |
| 22 | 43443 | 37715 | 37280 | 37040 | 36946 | 25525 |
| 23 | 45200 | 39358 | 38501 | 38267 | 38150 | 7002 |
| 25 | 44876 | 38282 | 37815 | 37624 | 37606 | 1076 |
| 27 | 39451 | 33895 | 33425 | 33339 | 33339 | 301 |
| 29 | 51098 | 44739 | 44318 | 44155 | 43907 | 22063 |
| 30 | 46917 | 39070 | 38828 | 38746 | 38746 | 1048 |
| 31 | 47137 | 41739 | 41522 | 40646 | 39447 | 39392 |
| 32 | 46971 | 40700 | 40231 | 39588 | 38857 | 38816 |
| 33 | 39391 | 34310 | 33848 | 33063 | 32052 | 32030 |
| 34 | 44566 | 34604 | 34197 | 33880 | 30418 | 23842 |
| 35 | 40681 | 35905 | 35698 | 35093 | 34171 | 34109 |
| 36 | 46321 | 39823 | 39461 | 39051 | 37221 | 21217 |
| 37 | 45570 | 39254 | 38858 | 37195 | 33997 | 33969 |
| 38 | 48423 | 42685 | 42236 | 41575 | 40666 | 40641 |
| 39 | 45562 | 38943 | 38562 | 38016 | 36846 | 24525 |
| 40 | 44638 | 39576 | 39198 | 36818 | 35059 | 35024 |
| 41 | 49912 | 31440 | 31190 | 31017 | 30950 | 7091 |
| 42 | 51752 | 43669 | 42635 | 41902 | 41902 | 1101 |
| 43 | 45004 | 38901 | 38480 | 38444 | 38444 | 1251 |
| 44 | 47440 | 40919 | 40478 | 39940 | 37780 | 27910 |
| 45 | 49568 | 43038 | 42635 | 42488 | 41530 | 24923 |
| 46 | 48670 | 33013 | 32793 | 32604 | 32546 | 22057 |
| 47 | 49955 | 43082 | 42833 | 42760 | 42754 | 781 |
| 48 | 48724 | 40383 | 40118 | 40069 | 40065 | 525 |
| 50 | 50830 | 43186 | 42186 | 41618 | 41577 | 8393 |
| 51 | 44203 | 38835 | 38555 | 38288 | 38164 | 38092 |
| 52 | 42192 | 37592 | 37299 | 36658 | 35360 | 34895 |
| 54 | 39930 | 34486 | 33066 | 30905 | 30748 | 20464 |
| 55 | 45851 | 40565 | 40144 | 39779 | 39433 | 39355 |
| 56 | 50436 | 43777 | 43498 | 43305 | 43282 | 43232 |
| 57 | 43569 | 38225 | 36808 | 34592 | 34489 | 32804 |
| 59 | 50085 | 42170 | 41790 | 40856 | 40049 | 39655 |
| 60 | 48445 | 41342 | 40456 | 39037 | 38848 | 38405 |
| 61 | 48781 | 42561 | 42331 | 41545 | 36929 | 36816 |
| 62 | 44796 | 38336 | 38052 | 37523 | 36705 | 36272 |
| 63 | 43288 | 38129 | 37385 | 36759 | 35866 | 35695 |
| 65 | 45751 | 40704 | 40188 | 39024 | 36079 | 36017 |
| 66 | 49821 | 44186 | 43831 | 43600 | 42938 | 42861 |
| 68 | 39343 | 34586 | 34173 | 33480 | 30232 | 19422 |
| 69 | 49292 | 42808 | 42115 | 41181 | 30909 | 30197 |
| 70 | 46895 | 40510 | 40003 | 38296 | 36365 | 36330 |
| 71 | 31754 | 27846 | 27435 | 26792 | 26072 | 26042 |
| 73 | 44442 | 38412 | 37958 | 37144 | 35345 | 35309 |
| 74 | 46866 | 40890 | 40360 | 40013 | 39668 | 20894 |
| 75 | 45549 | 39857 | 39555 | 38928 | 37644 | 23907 |
| 76 | 46635 | 40710 | 40079 | 39854 | 39854 | 1362 |
| 77 | 44654 | 38942 | 38372 | 38205 | 38121 | 10354 |
| 78 | 50244 | 43600 | 43204 | 43086 | 43086 | 438 |
| 79 | 44374 | 38990 | 38594 | 38449 | 37891 | 24309 |
| 80 | 47819 | 42155 | 41756 | 41332 | 32758 | 12705 |
| 81 | 43707 | 38240 | 37887 | 37056 | 33483 | 26409 |
| 82 | 45856 | 39226 | 38876 | 38644 | 38526 | 29092 |
| 84 | 47412 | 41244 | 40824 | 40503 | 39548 | 22411 |
| 85 | 53577 | 45928 | 45564 | 45115 | 41224 | 27219 |
| 86 | 42259 | 36878 | 36333 | 36122 | 35884 | 17801 |
| 88 | 43549 | 38026 | 37362 | 37161 | 37007 | 20806 |
| 89 | 44911 | 38399 | 37733 | 37413 | 37413 | 405 |
| 90 | 44730 | 38251 | 37585 | 37123 | 36172 | 20555 |
| 91 | 39013 | 33692 | 33047 | 31357 | 31262 | 19252 |
| 92 | 50113 | 43733 | 42969 | 42695 | 42689 | 2267 |
| 93 | 43991 | 38099 | 37288 | 36832 | 36818 | 3572 |
| 94 | 46503 | 40591 | 39957 | 39735 | 39524 | 11651 |
| 95 | 51320 | 44922 | 44164 | 43604 | 43483 | 11861 |
| 96 | 41715 | 35756 | 35114 | 34699 | 34390 | 19463 |
| 97 | 47550 | 39881 | 38266 | 37783 | 37677 | 14071 |
| 98 | 46453 | 40263 | 39645 | 39188 | 37552 | 27013 |
| 99 | 54559 | 46625 | 45112 | 44142 | 44101 | 12475 |
| 101 | 48077 | 42166 | 41635 | 41004 | 40680 | 39060 |
| 102 | 46328 | 38172 | 37462 | 36322 | 31986 | 31334 |
| 103 | 43495 | 38125 | 37589 | 37274 | 36981 | 32178 |
| 105 | 47789 | 40888 | 40297 | 39983 | 39629 | 20464 |
| 106 | 46944 | 40197 | 39407 | 38991 | 38814 | 9190 |
| 107 | 41401 | 36097 | 35620 | 35057 | 30774 | 26352 |
| 108 | 45969 | 39162 | 38296 | 37933 | 37921 | 665 |
| 109 | 51436 | 42621 | 42004 | 41635 | 41631 | 270 |
| 110 | 45681 | 39680 | 38873 | 38691 | 38648 | 8964 |
| 111 | 50786 | 43071 | 41813 | 41096 | 41075 | 5164 |
| 112.1 | 43741 | 36759 | 36199 | 35817 | 35765 | 24222 |
| 112.2 | 44542 | 38821 | 38306 | 38026 | 37935 | 23433 |
| 113 | 44414 | 38306 | 37671 | 37207 | 37023 | 24734 |
| 114 | 44720 | 37720 | 36985 | 36523 | 36085 | 14062 |
| 115 | 52244 | 44602 | 43644 | 42994 | 42874 | 9671 |
| 117 | 45730 | 38508 | 37320 | 36381 | 36378 | 1290 |
| 118 | 37358 | 31910 | 31620 | 31429 | 30828 | 28324 |
| 119 | 49695 | 41850 | 41045 | 40326 | 40317 | 3261 |
| 122 | 45211 | 39158 | 38451 | 38011 | 38011 | 842 |
| 123 | 52817 | 45764 | 44978 | 44447 | 44287 | 26690 |
| 124 | 40093 | 34335 | 33118 | 32329 | 32327 | 908 |
| 125 | 45075 | 36705 | 36383 | 35783 | 35729 | 18412 |
| 126 | 51146 | 43811 | 43221 | 40568 | 35064 | 35003 |
| 127 | 51384 | 44820 | 44497 | 43486 | 40434 | 40332 |
| 128 | 42207 | 36909 | 36450 | 35662 | 34536 | 34511 |
| 129 | 47194 | 40949 | 40440 | 39936 | 39087 | 39042 |
| 130 | 52668 | 41209 | 40828 | 39432 | 36072 | 36051 |
| 133 | 45414 | 39305 | 38620 | 37720 | 34968 | 34917 |
| 134 | 50417 | 43512 | 42819 | 41918 | 40924 | 40641 |
| 135 | 43641 | 35911 | 35516 | 34953 | 33397 | 32995 |
| 136 | 50883 | 43658 | 43258 | 42345 | 35746 | 35719 |
| 137 | 46616 | 40471 | 39862 | 39300 | 28056 | 27896 |
| 138 | 44478 | 37389 | 36860 | 36243 | 34652 | 34594 |
| 139 | 40807 | 27754 | 27322 | 26387 | 20292 | 20259 |
| 140 | 43677 | 38569 | 38164 | 37600 | 37497 | 37445 |
| 141 | 43671 | 38326 | 37920 | 37465 | 36412 | 36376 |
| 145 | 49064 | 40828 | 40323 | 39364 | 33978 | 33905 |
| 147 | 56180 | 47475 | 46801 | 46134 | 38718 | 38539 |
| 148.1 | 49103 | 42539 | 42039 | 40885 | 40775 | 40588 |
| 148.2 | 43886 | 38124 | 37785 | 37344 | 36480 | 36165 |
| 150 | 44608 | 38567 | 37708 | 36958 | 36918 | 16629 |
| 154 | 46120 | 40444 | 40193 | 39551 | 31232 | 31199 |
| 155 | 43740 | 37972 | 37301 | 37028 | 37003 | 8135 |
| 156 | 41332 | 35828 | 35147 | 34683 | 34276 | 18412 |
| 157 | 44730 | 38564 | 37678 | 37174 | 37078 | 11788 |
| 159 | 43639 | 38312 | 37959 | 37353 | 35190 | 34207 |
| 160 | 49241 | 41677 | 41139 | 40863 | 40863 | 925 |
| 161 | 47824 | 37820 | 36959 | 36276 | 35962 | 1515 |
| 162 | 47750 | 36092 | 35811 | 35010 | 31887 | 31735 |
| 163 | 45079 | 37410 | 37019 | 36798 | 36364 | 33483 |
| 164 | 49590 | 41561 | 40863 | 40332 | 40235 | 13758 |
| 166.1 | 43593 | 37847 | 37678 | 37437 | 37226 | 7857 |
| 166.2 | 52656 | 44462 | 44217 | 43526 | 42607 | 23821 |
| 167 | 42478 | 37148 | 36896 | 36761 | 36719 | 35168 |
| 169 | 44339 | 37835 | 37425 | 37012 | 36977 | 24471 |
| 171 | 40182 | 34832 | 34499 | 34285 | 33876 | 29983 |
| 173 | 43850 | 38279 | 37788 | 37430 | 37093 | 16566 |
| 174 | 38214 | 32522 | 32344 | 31596 | 30507 | 23518 |
| 175 | 41812 | 36830 | 36593 | 34541 | 31252 | 28972 |
| 176 | 47933 | 41148 | 40339 | 40000 | 40000 | 444 |
| 177 | 52306 | 45296 | 44809 | 44570 | 44563 | 405 |
| 178 | 37588 | 31983 | 31224 | 30705 | 30705 | 324 |
| 180.1 | 47459 | 39894 | 39304 | 38838 | 38421 | 28477 |
| 180.2 | 43208 | 36790 | 36449 | 36093 | 35093 | 23952 |
| 181.1 | 48528 | 41541 | 40859 | 40304 | 40221 | 16642 |
| 181.2 | 50046 | 43517 | 43180 | 43048 | 42870 | 21696 |
| 182.1 | 44878 | 38005 | 37198 | 36786 | 36566 | 20232 |
| 182.2 | 37747 | 31964 | 31475 | 31220 | 30767 | 16694 |
| 183 | 41270 | 36172 | 35701 | 35309 | 34029 | 16970 |
| 184 | 37139 | 32145 | 31668 | 31451 | 31326 | 14368 |
| 187 | 43567 | 37083 | 36433 | 35976 | 35528 | 16101 |
| 188 | 51525 | 44506 | 44129 | 43815 | 42922 | 32702 |
| 189 | 48190 | 40720 | 40159 | 39970 | 39779 | 17364 |
| 190 | 46438 | 39946 | 39375 | 38872 | 37024 | 33937 |
| 191 | 46589 | 37147 | 36697 | 36409 | 33903 | 28965 |
| 192 | 47307 | 40829 | 40513 | 40128 | 38448 | 27531 |
| 193 | 48977 | 42834 | 42024 | 41221 | 38792 | 38725 |
| 194 | 45386 | 39771 | 39476 | 39031 | 36820 | 36780 |
| 195 | 41675 | 36346 | 35751 | 34982 | 34143 | 34088 |
| 196 | 47388 | 35080 | 34289 | 33112 | 31689 | 31632 |
| 197 | 42460 | 37466 | 37123 | 36405 | 34997 | 34976 |
| 198 | 44378 | 38644 | 38286 | 37724 | 36367 | 36339 |
| 199 | 40485 | 35522 | 35152 | 34741 | 33618 | 33574 |
| 200 | 48351 | 40013 | 39511 | 38721 | 32645 | 32607 |
| 201.1 | 44573 | 38974 | 38424 | 37906 | 37021 | 36961 |
| 201.2 | 42668 | 37384 | 36947 | 36275 | 33593 | 33537 |
| 202 | 43874 | 38765 | 38522 | 37667 | 37366 | 37339 |
| 203 | 47463 | 40335 | 40070 | 39529 | 36837 | 36791 |
| 204 | 41814 | 35943 | 35469 | 34705 | 32730 | 32688 |
| 205 | 45974 | 39253 | 38520 | 37883 | 36991 | 36724 |
| 206 | 44136 | 37451 | 36519 | 35627 | 35607 | 4099 |
| 207 | 50670 | 44006 | 43555 | 43087 | 42819 | 42791 |
| 208 | 48783 | 41532 | 41005 | 40494 | 39615 | 39524 |
| 209 | 48712 | 42575 | 42262 | 41039 | 37045 | 36988 |
| 210 | 42566 | 36264 | 35826 | 35346 | 34005 | 33932 |
| 212 | 38607 | 33139 | 32491 | 32113 | 31190 | 31164 |
| 213 | 42060 | 36675 | 36162 | 35223 | 34414 | 34381 |
| 214 | 34256 | 30075 | 29420 | 28839 | 27999 | 27888 |
| 215 | 38823 | 34032 | 33734 | 33234 | 31662 | 31640 |
| 216 | 46186 | 39238 | 38682 | 37807 | 36881 | 36825 |
| 217 | 47337 | 40201 | 39680 | 39250 | 36683 | 36646 |
| 218 | 44411 | 36697 | 35376 | 34599 | 32563 | 32518 |
| 219 | 41128 | 36054 | 35511 | 35260 | 34153 | 34106 |
| 220 | 41452 | 35715 | 34622 | 34149 | 32491 | 32433 |
| 221 | 42627 | 37118 | 36547 | 36028 | 32371 | 32345 |
| 222 | 42609 | 37290 | 36785 | 36075 | 31849 | 31725 |
| 223 | 44698 | 37339 | 37001 | 36463 | 32287 | 32257 |
| 224 | 38112 | 32953 | 31258 | 29806 | 29576 | 28989 |
| 225 | 46840 | 39642 | 39089 | 38574 | 35174 | 35153 |
| 226 | 49549 | 44090 | 43777 | 43558 | 43398 | 43343 |
| 227 | 45191 | 38909 | 38416 | 38087 | 37603 | 25482 |
| 228 | 44435 | 37774 | 36116 | 35323 | 35192 | 17642 |
| 229 | 47424 | 40367 | 40028 | 39852 | 37690 | 26458 |
| 230 | 52393 | 43602 | 42010 | 40432 | 40390 | 2755 |
| 231 | 43175 | 36711 | 35992 | 35672 | 35480 | 18611 |
| 232 | 45648 | 39407 | 38579 | 38175 | 38053 | 9519 |
| 233 | 41399 | 36102 | 35605 | 35401 | 35284 | 24832 |
| 234 | 46350 | 39822 | 39259 | 38800 | 38487 | 11875 |
| 235 | 50002 | 42962 | 42411 | 41078 | 40893 | 15844 |
| 236 | 43197 | 37020 | 35899 | 35513 | 34694 | 21032 |
| 237 | 48954 | 42387 | 41565 | 41224 | 41224 | 925 |
| 238 | 46228 | 39379 | 37908 | 37270 | 37270 | 880 |
| 239 | 49229 | 41155 | 40564 | 40386 | 40375 | 10737 |
| 240 | 42737 | 36788 | 35520 | 34475 | 34468 | 7033 |
| 241 | 45485 | 38970 | 38210 | 38102 | 37933 | 25665 |
| 242 | 46050 | 40614 | 40038 | 39035 | 38769 | 38719 |
| 244.1 | 51860 | 43592 | 43155 | 42591 | 41704 | 41615 |
| 244.2 | 44004 | 38336 | 38133 | 37583 | 35602 | 35585 |
| 245.1 | 37030 | 32255 | 30943 | 29287 | 28893 | 28869 |
| 245.2 | 45619 | 36626 | 35770 | 34245 | 33017 | 32915 |
| 246.1 | 45041 | 39087 | 38177 | 37349 | 37131 | 37057 |
| 246.2 | 47210 | 41236 | 40814 | 40404 | 39636 | 39583 |
| 247 | 42424 | 37158 | 36959 | 36516 | 34214 | 34188 |
| 248 | 46723 | 39570 | 39017 | 38328 | 37763 | 37724 |
| 249.1 | 43478 | 37515 | 36797 | 35920 | 35619 | 35564 |
| 249.2 | 49713 | 42805 | 42474 | 41252 | 39658 | 39579 |
| 250 | 50813 | 38070 | 36989 | 36311 | 33355 | 33274 |
| 251.1 | 45377 | 38116 | 37524 | 36898 | 36580 | 36539 |
| 251.2 | 43282 | 37554 | 37140 | 36591 | 35304 | 35275 |
| 272 | 1481 | 1353 | 1337 | 1282 | 1282 | 1236 |
| 273 | 52373 | 50482 | 50129 | 49551 | 46856 | 46771 |
| 275 | 787 | 730 | 723 | 666 | 663 | 622 |
| 276 | 29877 | 28984 | 28816 | 28536 | 28041 | 28007 |
| 277 | 699 | 619 | 580 | 555 | 549 | 530 |
| 278 | 1548 | 1442 | 1415 | 1349 | 1349 | 1337 |
| 279 | 1450 | 1343 | 1299 | 1295 | 1295 | 1289 |
| 280 | 10318 | 9594 | 9145 | 8414 | 8406 | 8301 |
| 283 | 381 | 329 | 319 | 311 | 311 | 223 |
| 284 | 60977 | 58327 | 58099 | 58043 | 58043 | 3045 |
| 285 | 57769 | 55716 | 55637 | 55067 | 54891 | 35639 |
| 286 | 49132 | 47211 | 47090 | 46955 | 46924 | 21388 |
| 287 | 70164 | 67991 | 67821 | 67686 | 64858 | 54924 |
| 288 | 38700 | 36299 | 35807 | 35697 | 35697 | 16204 |
| 289 | 43946 | 42345 | 42313 | 42297 | 42297 | 35989 |
| 296 | 61434 | 59486 | 59342 | 58656 | 55475 | 55450 |
| 297 | 3252 | 2999 | 2924 | 2840 | 2838 | 2804 |
| 298 | 66861 | 64486 | 64104 | 63499 | 62956 | 62889 |
| 300 | 3925 | 3642 | 3548 | 3522 | 3522 | 3475 |
| 306 | 1307 | 1194 | 1160 | 1157 | 1157 | 582 |
| 307 | 82065 | 78491 | 78173 | 78044 | 77982 | 21821 |
| 308 | 67388 | 64724 | 64423 | 64262 | 64156 | 26996 |
| 309 | 73619 | 70527 | 70176 | 70021 | 70021 | 10296 |
| 310 | 4501 | 4120 | 4076 | 4059 | 4059 | 1349 |
| 311 | 85496 | 82178 | 81795 | 81657 | 81657 | 12118 |
| 312 | 68671 | 61119 | 59885 | 59068 | 59068 | 2046 |
| 313 | 6433 | 6061 | 6045 | 6041 | 6041 | 193 |
| 322 | 78834 | 76830 | 76637 | 75892 | 72292 | 72242 |
| 324 | 66270 | 64436 | 64203 | 63701 | 59716 | 59643 |
| 325 | 78371 | 75469 | 75227 | 74368 | 70022 | 70000 |
| 326 | 3713 | 3487 | 3448 | 3423 | 3423 | 3410 |
| 327 | 62038 | 60254 | 60004 | 59379 | 57822 | 57785 |
| 328 | 77436 | 74416 | 74117 | 72903 | 67213 | 67119 |
| 330 | 76249 | 72782 | 71964 | 70559 | 70471 | 29574 |
| 334 | 92827 | 89541 | 89116 | 89002 | 88710 | 46843 |
| 335 | 67489 | 61362 | 60636 | 60124 | 60122 | 22325 |
| 336 | 73592 | 71111 | 70781 | 70463 | 69540 | 56905 |
| 337 | 75200 | 70391 | 69988 | 69886 | 69886 | 1367 |
| 339 | 37658 | 35399 | 34962 | 34758 | 34755 | 9789 |
| 344 | 80904 | 78420 | 78166 | 76811 | 63364 | 63227 |
| 345 | 64537 | 62261 | 61969 | 61743 | 59851 | 59797 |
| 346 | 83453 | 80315 | 80060 | 78134 | 69659 | 69513 |
| 347 | 71735 | 69354 | 69141 | 69017 | 69017 | 68998 |
| 349 | 89038 | 85659 | 85351 | 83482 | 76827 | 76812 |
| 350 | 63833 | 61697 | 61393 | 60273 | 55929 | 55662 |
| 354 | 19215 | 18009 | 17916 | 17908 | 17908 | 7097 |
| 355 | 42957 | 36531 | 35170 | 34742 | 34742 | 670 |
| 356 | 119182 | 107586 | 106645 | 106295 | 105913 | 32978 |
| 357 | 98388 | 93743 | 93348 | 93099 | 92096 | 61778 |
| 360 | 91175 | 86795 | 86291 | 86083 | 85949 | 30376 |
| 362 | 102840 | 97740 | 97304 | 96533 | 93769 | 52226 |
| 364 | 79215 | 75792 | 75052 | 74208 | 71009 | 70982 |
| 365 | 92270 | 88515 | 88019 | 86909 | 76867 | 76834 |
| 368 | 53455 | 50827 | 50478 | 50083 | 46638 | 46595 |
| 369 | 82647 | 78846 | 78112 | 77030 | 75004 | 74915 |
| 370 | 77881 | 71908 | 70977 | 70266 | 65762 | 65724 |
| 374 | 83517 | 78579 | 78247 | 78129 | 78127 | 952 |
| 376 | 97475 | 92839 | 92457 | 92391 | 92371 | 45581 |
| 377 | 87225 | 83447 | 83232 | 83155 | 82979 | 43111 |
| 378 | 88004 | 81473 | 80821 | 80647 | 80340 | 22774 |
| 380 | 82291 | 77779 | 77506 | 77147 | 76476 | 52083 |
| 381 | 65580 | 62358 | 61929 | 61713 | 61509 | 18528 |
| 382 | 67203 | 63254 | 62604 | 62449 | 62449 | 599 |
| 384 | 64816 | 62487 | 62358 | 61625 | 53627 | 53540 |
| 385 | 67301 | 65233 | 64991 | 64655 | 62870 | 62585 |
| 387 | 79843 | 77102 | 76866 | 75795 | 71954 | 71872 |
| 388 | 1816 | 1483 | 1473 | 1430 | 1414 | 1407 |
| 393 | 71269 | 66889 | 65790 | 63863 | 60847 | 60514 |
| 394 | 74074 | 69846 | 69554 | 69501 | 69501 | 276 |
| 395 | 81497 | 77144 | 76886 | 76771 | 76624 | 11219 |
| 396 | 1091 | 975 | 952 | 945 | 944 | 59 |
| 397 | 81857 | 78006 | 77760 | 77723 | 77723 | 590 |
| 398 | 90341 | 83922 | 82872 | 82496 | 82496 | 4253 |
| 399 | 44429 | 42060 | 41796 | 41754 | 41754 | 15032 |
| 400 | 46955 | 44630 | 44481 | 44382 | 44364 | 6872 |
| 405 | 78849 | 75794 | 75279 | 74518 | 72594 | 63519 |
| 406 | 82509 | 78379 | 77760 | 77467 | 77467 | 555 |
| 408 | 83841 | 80102 | 79432 | 77548 | 74370 | 72751 |
| 409 | 80299 | 77661 | 77034 | 76789 | 76481 | 76433 |
| 410 | 89083 | 84489 | 83907 | 83724 | 83711 | 3227 |
| 414 | 96759 | 92050 | 91732 | 91664 | 91664 | 183 |
| 415 | 88419 | 84883 | 84692 | 84622 | 84592 | 8949 |
| 416 | 93211 | 88537 | 88221 | 88105 | 88105 | 461 |
| 417 | 99605 | 95782 | 95633 | 95308 | 91649 | 53628 |
| 418 | 82286 | 76096 | 75386 | 75290 | 75244 | 1062 |
| 420 | 19616 | 17671 | 17189 | 17079 | 17079 | 2390 |
| 422 | 18798 | 17530 | 17362 | 17339 | 17339 | 277 |
| 424 | 1931 | 1730 | 1666 | 1653 | 1653 | 360 |
| 425 | 17765 | 16922 | 16840 | 16817 | 16789 | 10096 |
| 426 | 89727 | 82789 | 81709 | 80630 | 78760 | 78386 |
| 428 | 110591 | 103120 | 101670 | 99552 | 96656 | 93833 |
| 430 | 68518 | 65237 | 64758 | 64421 | 61668 | 61632 |
| 431 | 91690 | 86312 | 85294 | 84462 | 81540 | 57106 |
| 434 | 63461 | 58910 | 58307 | 58157 | 57943 | 8230 |
| 435 | 71310 | 66829 | 66568 | 66442 | 65284 | 30876 |
| 437 | 79909 | 72877 | 72001 | 71780 | 71752 | 2755 |
| 438 | 81768 | 76135 | 75729 | 75540 | 74464 | 23907 |
| 440 | 40939 | 36343 | 36134 | 36100 | 35994 | 2662 |
| 442 | 90498 | 83004 | 82097 | 81499 | 81251 | 37263 |
| 443 | 42809 | 36665 | 35396 | 34180 | 34180 | 1560 |
| 444 | 74560 | 71468 | 71257 | 70454 | 67902 | 67863 |
| 445 | 77100 | 66659 | 65451 | 62592 | 61802 | 61351 |
| 446 | 65936 | 62734 | 62540 | 62471 | 62151 | 13952 |
| 447 | 85263 | 81954 | 81640 | 80140 | 76424 | 76322 |
| 448 | 69855 | 67104 | 66199 | 64807 | 59018 | 58766 |
| 449 | 91014 | 85919 | 85428 | 84870 | 84438 | 42366 |
| 450 | 70059 | 65056 | 64080 | 63111 | 61534 | 61347 |
| 454 | 34065 | 32389 | 31962 | 31702 | 31702 | 1547 |
| 455 | 78660 | 75459 | 75239 | 75147 | 75147 | 38114 |
| 458 | 75741 | 72734 | 72614 | 72553 | 72430 | 22250 |
| 459 | 72601 | 68908 | 67590 | 66970 | 66946 | 13494 |
| 460 | 82634 | 79292 | 78827 | 78693 | 78435 | 22231 |
| 462 | 93508 | 90053 | 89947 | 89686 | 88567 | 32530 |
| 463 | 68742 | 65460 | 65203 | 65108 | 65066 | 21090 |
| 464 | 72110 | 70195 | 70071 | 69879 | 69000 | 68882 |
| 465 | 4577 | 3644 | 3379 | 2963 | 2956 | 2792 |
| 466 | 80290 | 77661 | 77558 | 77409 | 77038 | 76986 |
| 467 | 5710 | 5245 | 5164 | 5126 | 5125 | 5099 |
| 468 | 61512 | 58976 | 58597 | 58189 | 57056 | 57006 |
| 469 | 7349 | 6453 | 6168 | 5935 | 633 | 501 |
| 470 | 67906 | 65721 | 65578 | 65410 | 65260 | 64613 |
| 474 | 81204 | 78235 | 78027 | 77901 | 77566 | 42523 |
| 475 | 78867 | 74801 | 74420 | 74363 | 74363 | 921 |
| 476 | 53117 | 50527 | 50369 | 50281 | 50251 | 5652 |
| 477 | 66087 | 62934 | 62583 | 62450 | 62450 | 2485 |
| 479 | 73630 | 70617 | 70437 | 70339 | 70304 | 35234 |
| 480 | 59415 | 57117 | 56990 | 56858 | 56823 | 43812 |
| 483 | 72941 | 69584 | 69246 | 69180 | 69180 | 807 |
| 484 | 68052 | 65879 | 65668 | 64731 | 63084 | 63026 |
| 485 | 69451 | 66895 | 66582 | 65972 | 65252 | 65117 |
| 486 | 87445 | 84550 | 84192 | 82579 | 70908 | 70790 |
| 487 | 32071 | 31009 | 30845 | 30691 | 30546 | 30509 |
| 489 | 51231 | 49420 | 49150 | 48535 | 48197 | 48141 |
| 490 | 57341 | 55232 | 54765 | 53929 | 41862 | 24262 |
| 494 | 42433 | 40525 | 40258 | 40118 | 39998 | 22129 |
| 495 | 69135 | 67221 | 67169 | 67135 | 67107 | 46862 |
| 497 | 48975 | 47084 | 46944 | 46889 | 46871 | 3966 |
| 499 | 55172 | 53456 | 53307 | 53242 | 52806 | 43631 |
| 500 | 62133 | 59969 | 59812 | 59545 | 59536 | 37481 |
| 501 | 45 | 31 | 15 | 15 | 15 | 3 |
| 502 | 65419 | 62691 | 62526 | 62450 | 62372 | 18700 |
| 505 | 61354 | 59188 | 58834 | 58039 | 54371 | 54171 |
| 508 | 65984 | 63601 | 63318 | 62110 | 50133 | 50060 |
| 509 | 65414 | 63479 | 63143 | 62427 | 60102 | 60046 |
| 511 | 78258 | 76092 | 75843 | 75698 | 74670 | 74603 |
| 514 | 73246 | 70991 | 70616 | 70202 | 68898 | 68823 |
| 517 | 66859 | 64686 | 64368 | 64072 | 62844 | 62324 |
| 522 | 13525 | 12831 | 12563 | 12336 | 12335 | 12174 |
| 527 | 45278 | 43500 | 43315 | 43230 | 43230 | 370 |
| 528 | 55842 | 53564 | 53460 | 53407 | 53407 | 13921 |
| 529 | 82957 | 78566 | 78116 | 77966 | 77683 | 3931 |
| 530 | 87301 | 82973 | 82541 | 82403 | 82062 | 18652 |
| 531 | 72521 | 69159 | 68935 | 68871 | 68430 | 1140 |
| 532 | 63540 | 60113 | 59768 | 59566 | 59566 | 507 |
| 533 | 79226 | 75507 | 75091 | 74854 | 74577 | 18854 |
| 536 | 68726 | 65994 | 65863 | 65801 | 65801 | 4657 |
| 537 | 6016 | 5632 | 5466 | 5404 | 5404 | 1031 |
| 538 | 77160 | 73826 | 73265 | 73111 | 73069 | 15945 |
| 539 | 81022 | 77849 | 77684 | 77463 | 76454 | 34074 |
| 544 | 57928 | 55424 | 55061 | 54889 | 54889 | 2324 |
| 545 | 53309 | 51327 | 51280 | 51224 | 51224 | 51 |
| 546 | 80845 | 76594 | 76153 | 75947 | 75932 | 2921 |
| 547 | 14250 | 12055 | 11285 | 10059 | 10001 | 7953 |
| 548 | 3898 | 2290 | 1945 | 1464 | 1453 | 441 |
| 549 | 83908 | 80166 | 79079 | 77283 | 75842 | 75458 |
| 551 | 83856 | 79521 | 79013 | 78712 | 78709 | 322 |
| 552 | 73922 | 71481 | 70976 | 70456 | 70385 | 70050 |
| 553 | 11427 | 9241 | 8630 | 7701 | 7666 | 7077 |
| 554 | 67903 | 64942 | 64673 | 64079 | 60230 | 59830 |
| 555 | 69170 | 66654 | 65743 | 64542 | 56903 | 56770 |
| 556 | 58261 | 53644 | 52201 | 45144 | 44645 | 40194 |
| 557 | 67176 | 64928 | 64637 | 64288 | 61958 | 61866 |
| 558 | 68900 | 65496 | 64447 | 61454 | 57331 | 54738 |
| 559 | 57614 | 52821 | 51495 | 47438 | 47248 | 46070 |
| 560 | 71146 | 68083 | 67399 | 65601 | 62721 | 62376 |
| 561 | 62917 | 60438 | 59705 | 57950 | 57714 | 57032 |
| 567 | 67245 | 63682 | 63088 | 62786 | 62786 | 4443 |
| 568 | 52969 | 50921 | 50868 | 50822 | 50816 | 12622 |
| 569 | 52721 | 49653 | 49312 | 49148 | 48962 | 8169 |
| 570 | 77629 | 74228 | 73923 | 73768 | 73459 | 53492 |
| 571 | 75233 | 71507 | 71008 | 70677 | 70542 | 7022 |
| 572 | 83262 | 79616 | 79387 | 79277 | 79269 | 2949 |
| 573 | 19122 | 18077 | 17991 | 17962 | 17962 | 141 |
| 590 | 69198 | 66421 | 66274 | 66217 | 66123 | 27111 |
| 591 | 28426 | 26722 | 26391 | 26169 | 26022 | 14326 |
| 592 | 60158 | 57301 | 56980 | 56872 | 56872 | 2304 |
| 593 | 66753 | 64240 | 64033 | 63946 | 63807 | 35609 |
| 594 | 80493 | 76496 | 75978 | 75735 | 75660 | 17810 |
| 595 | 79447 | 76883 | 76675 | 76523 | 75114 | 56824 |
| 596 | 7141 | 6420 | 6352 | 6339 | 6339 | 2917 |
| 600 | 25322 | 24081 | 23940 | 23725 | 22754 | 22726 |
| 602 | 13801 | 12983 | 12886 | 12834 | 12802 | 12772 |
| 603 | 77156 | 74115 | 73621 | 73029 | 72170 | 71975 |
| 604 | 49091 | 45668 | 45138 | 44018 | 43946 | 42708 |
| 605 | 67446 | 65030 | 64785 | 64534 | 64066 | 63986 |
| 606 | 55361 | 53471 | 53137 | 52770 | 51852 | 51783 |
| 612 | 65975 | 60003 | 59524 | 59385 | 59250 | 23380 |
| 613 | 78955 | 74125 | 73750 | 73667 | 73648 | 1011 |
| 614 | 86448 | 78227 | 77776 | 77674 | 77674 | 26187 |
| 615 | 73267 | 67288 | 66797 | 66575 | 66575 | 1357 |
| 616 | 60720 | 53127 | 52342 | 52196 | 52196 | 7498 |
| 617 | 70135 | 66234 | 65949 | 65323 | 62990 | 47909 |
| 618 | 76714 | 73353 | 73108 | 72829 | 71378 | 66724 |
| 619 | 81155 | 73627 | 73199 | 72183 | 61220 | 61059 |
| 620 | 77292 | 74088 | 73889 | 72672 | 69968 | 69906 |
| 621 | 65301 | 55981 | 54249 | 52371 | 52097 | 52010 |
| 622 | 66268 | 64026 | 63795 | 63110 | 61695 | 61601 |
| 623 | 61760 | 52410 | 50253 | 40863 | 40583 | 10817 |
| 624 | 57851 | 47142 | 45523 | 43035 | 42879 | 42821 |
| 625 | 78560 | 74429 | 73814 | 72349 | 61132 | 61049 |
| 627 | 67412 | 64435 | 64302 | 64213 | 64195 | 22273 |
| 628 | 74833 | 70886 | 70522 | 70228 | 69666 | 36433 |
| 629 | 56494 | 51986 | 51601 | 51507 | 51507 | 626 |
| 633 | 55716 | 52640 | 52180 | 51903 | 51264 | 29669 |
| 634 | 62911 | 59274 | 58925 | 58859 | 58695 | 22199 |
| 635 | 77010 | 74020 | 73730 | 73507 | 72806 | 61313 |
| 636 | 93337 | 89411 | 89018 | 88719 | 86100 | 28013 |
| 637 | 71168 | 69071 | 68867 | 68640 | 64354 | 64311 |
| 638 | 48787 | 47061 | 46938 | 46129 | 39740 | 39713 |
| 639 | 1356 | 1017 | 805 | 529 | 529 | 434 |
| 641 | 2726 | 2433 | 2257 | 2102 | 2099 | 1987 |
| 642 | 49719 | 48324 | 47882 | 47322 | 47135 | 47055 |
| 643 | 67616 | 64981 | 64107 | 61723 | 60702 | 59959 |
| 656 | 80326 | 73811 | 73607 | 73404 | 73231 | 73222 |
| 657 | 69662 | 63399 | 63240 | 62947 | 62195 | 60727 |
| 659 | 3111 | 2278 | 2106 | 1909 | 1909 | 1523 |
| 660 | 70634 | 63108 | 62853 | 62484 | 60682 | 59343 |
| 664 | 66670 | 60986 | 60922 | 60906 | 60900 | 60882 |
| 665 | 61256 | 54630 | 54518 | 53230 | 51303 | 51285 |
| 666 | 41311 | 36845 | 36793 | 36751 | 36658 | 16144 |
| 667 | 8934 | 7653 | 7590 | 7574 | 7574 | 787 |
| 668 | 58408 | 52184 | 52109 | 52040 | 52040 | 13401 |
| 670 | 80397 | 73193 | 73071 | 72619 | 71894 | 53709 |
| 671 | 65363 | 58654 | 58493 | 58454 | 58382 | 23493 |
| 672 | 31478 | 27822 | 27783 | 27767 | 27767 | 138 |
| 674 | 26619 | 23982 | 23967 | 23951 | 23887 | 11086 |
| 676 | 87676 | 78691 | 78470 | 78181 | 77563 | 53541 |
| 677 | 69703 | 62486 | 62365 | 62028 | 61996 | 23911 |
| 679 | 74989 | 67549 | 67455 | 67358 | 67163 | 44766 |
| 680 | 62377 | 55290 | 55229 | 55178 | 55104 | 4627 |
| 682 | 71613 | 65756 | 65525 | 65298 | 57202 | 57168 |
| 683 | 105713 | 95809 | 95539 | 95142 | 86430 | 86307 |
| 686 | 24703 | 22578 | 22418 | 22252 | 20224 | 20150 |
| 687 | 93245 | 84451 | 84281 | 83859 | 82196 | 82173 |
| 688 | 60215 | 54923 | 54639 | 54255 | 53043 | 53013 |
| 689 | 23805 | 21523 | 21400 | 21230 | 20749 | 20677 |
| 690 | 87794 | 80163 | 80022 | 79836 | 78745 | 78745 |
| 693 | 54084 | 49096 | 48782 | 48438 | 42982 | 42665 |
| 694 | 71240 | 65652 | 65497 | 64897 | 62713 | 62696 |
| 695 | 91229 | 84473 | 84339 | 84078 | 82048 | 81976 |
| 696 | 72699 | 66688 | 66305 | 65977 | 65566 | 65450 |
| 697 | 35187 | 31795 | 31636 | 31457 | 31440 | 31322 |
| 698 | 68549 | 62406 | 62332 | 61906 | 60939 | 60927 |
| 699 | 74841 | 67665 | 67537 | 67432 | 66561 | 45247 |
| 700 | 22055 | 20097 | 20074 | 20006 | 19702 | 19697 |
| 701 | 84865 | 77065 | 76796 | 76347 | 74042 | 73996 |
| 702 | 34590 | 30636 | 30558 | 30541 | 30541 | 5 |
| 704 | 59442 | 53456 | 53320 | 53265 | 53179 | 26732 |
| 705 | 51261 | 45022 | 44932 | 44865 | 44865 | 1232 |
| 706 | 47686 | 40895 | 40784 | 40693 | 40015 | 14953 |
| 707 | 75875 | 66942 | 66765 | 66726 | 66726 | 208 |
| 708 | 39345 | 34350 | 34191 | 34140 | 34140 | 421 |
| 709 | 30319 | 26933 | 26893 | 26881 | 26881 | 1804 |
| 710 | 9473 | 8418 | 8378 | 8374 | 8374 | 103 |
| 711 | 82086 | 75027 | 74906 | 74774 | 74352 | 67965 |
| 713 | 11011 | 9753 | 9737 | 9733 | 9733 | 1392 |
| 714 | 78444 | 71975 | 71836 | 71624 | 70392 | 59151 |
| 715 | 9589 | 8489 | 8472 | 8467 | 8467 | 71 |
| 716 | 81352 | 72167 | 71983 | 71723 | 71688 | 22884 |
| 718 | 39708 | 35827 | 35773 | 35729 | 8258 | 1002 |
| 720 | 1284 | 1111 | 1100 | 1097 | 1097 | 17 |
| 721 | 71787 | 63370 | 63280 | 63226 | 63226 | 333 |
| 722 | 91466 | 83691 | 83477 | 83201 | 82557 | 81977 |
| 723 | 100198 | 91374 | 91169 | 90233 | 87185 | 87158 |
| 724 | 83945 | 77162 | 76995 | 76627 | 73763 | 73753 |
| 726 | 74284 | 67885 | 67748 | 67514 | 66582 | 65025 |
| 727 | 24468 | 22601 | 22543 | 22374 | 21654 | 21645 |
| 728 | 16063 | 14762 | 14709 | 14677 | 14656 | 14634 |
| 729 | 99244 | 91159 | 90923 | 90428 | 89975 | 88418 |
| 730 | 77524 | 71483 | 71241 | 70871 | 70447 | 69995 |
| 732 | 9964 | 8870 | 8839 | 8810 | 8796 | 2526 |
| 733 | 14514 | 12978 | 12955 | 12943 | 12943 | 1931 |
| 734 | 24401 | 22019 | 21998 | 21955 | 21937 | 17764 |
| 735 | 43999 | 39735 | 39707 | 39659 | 39612 | 17886 |
| 736 | 9132 | 7974 | 7959 | 7957 | 7957 | 49 |
| 737 | 25612 | 22327 | 22259 | 22231 | 22231 | 75 |
| 738 | 7694 | 6651 | 6616 | 6613 | 6613 | 2018 |
| 740 | 73219 | 64795 | 64593 | 64479 | 64479 | 227 |
| 741 | 40701 | 35948 | 35896 | 35885 | 35885 | 2521 |
| 742 | 90933 | 82796 | 82599 | 82308 | 81570 | 81553 |
| 743 | 66170 | 61035 | 60835 | 60438 | 57210 | 57186 |
| 744 | 32009 | 29162 | 29075 | 28899 | 28656 | 28616 |
| 745.1 | 11382 | 10432 | 10395 | 10344 | 9847 | 9847 |
| 745.2 | 3712 | 3400 | 3379 | 3361 | 3192 | 3192 |
| 747 | 82390 | 75310 | 75154 | 74809 | 73293 | 73278 |
| 748 | 100861 | 92278 | 92039 | 91658 | 91014 | 91006 |
| 749 | 98182 | 89834 | 89699 | 89323 | 88502 | 88481 |
| 750 | 68124 | 62351 | 62123 | 61907 | 61777 | 61736 |
| 751 | 81921 | 73953 | 73803 | 73511 | 73033 | 54287 |
| 752 | 22695 | 20093 | 20066 | 20052 | 20052 | 1200 |
| 753 | 87443 | 77532 | 77475 | 77448 | 77440 | 6982 |
| 754 | 58000 | 52010 | 51954 | 51902 | 51871 | 8725 |
| 755 | 89926 | 81487 | 81334 | 80961 | 80627 | 45298 |
| 756 | 11277 | 10279 | 10258 | 10243 | 10228 | 10062 |
| 758 | 90373 | 80049 | 79806 | 79673 | 79673 | 203 |
| 759 | 70687 | 63211 | 62873 | 62732 | 62554 | 22554 |
| 760 | 24586 | 21923 | 21882 | 21855 | 21855 | 5237 |
| 761 | 49423 | 45069 | 45001 | 43975 | 43955 | 38844 |
| 764 | 5065 | 4411 | 4358 | 4298 | 4290 | 4152 |
| 766 | 41465 | 36844 | 36770 | 36622 | 33683 | 33660 |
| 767 | 14337 | 12829 | 12772 | 12618 | 12584 | 12495 |
| 769 | 80440 | 73159 | 73049 | 72618 | 72114 | 72050 |
| 770 | 4576 | 4091 | 4054 | 3998 | 3986 | 3947 |
| 771 | 107867 | 98719 | 98507 | 97696 | 96971 | 96951 |
| 772 | 97651 | 87159 | 87021 | 86162 | 83228 | 83206 |
| 774 | 96445 | 88204 | 88059 | 87243 | 54426 | 54366 |
| 776 | 82538 | 73686 | 73405 | 73124 | 72636 | 72630 |
| 777 | 55146 | 50853 | 50730 | 50507 | 46032 | 45996 |
| 778 | 73555 | 67152 | 67029 | 66789 | 66501 | 66470 |
| 779 | 81808 | 74260 | 74061 | 73339 | 68149 | 68123 |
| 780 | 47317 | 43205 | 43105 | 42966 | 42676 | 42646 |
| 781 | 36646 | 33341 | 33159 | 32758 | 32322 | 32292 |
| 783 | 44968 | 38595 | 38464 | 38425 | 38142 | 913 |
| 784 | 72708 | 64221 | 64121 | 64040 | 64040 | 1116 |
| 786 | 80175 | 71646 | 71573 | 71526 | 71526 | 120 |
| 787.1 | 43160 | 38206 | 38164 | 38152 | 38152 | 3271 |
| 787.2 | 13226 | 11744 | 11738 | 11731 | 11731 | 775 |
| 789 | 72434 | 65341 | 65166 | 65088 | 65051 | 41027 |
| 790 | 20514 | 17923 | 17877 | 17863 | 17863 | 219 |
| 791 | 28430 | 25376 | 25317 | 25285 | 25285 | 439 |
| 792 | 96926 | 89518 | 89340 | 88571 | 86219 | 86204 |
| 793 | 81089 | 74225 | 73924 | 73435 | 69495 | 69414 |
| 795 | 76540 | 69913 | 69653 | 69088 | 67011 | 66976 |
| 796 | 108701 | 99024 | 98895 | 96646 | 93231 | 93213 |
| 797 | 90947 | 83011 | 82860 | 82397 | 78852 | 78831 |
| 798 | 98135 | 89783 | 89674 | 89290 | 73394 | 73382 |
| 799 | 28885 | 26520 | 26384 | 26205 | 25425 | 25408 |
| 800 | 71329 | 63771 | 63525 | 62025 | 60782 | 57814 |
| 801 | 70971 | 65274 | 65159 | 65073 | 64626 | 64600 |
| 802 | 72542 | 64709 | 64614 | 64043 | 57643 | 21324 |
| 803 | 72997 | 65267 | 65139 | 65008 | 64930 | 39658 |
| 804 | 64928 | 58130 | 57996 | 57893 | 57556 | 22623 |
| 805 | 3260 | 2502 | 2407 | 2292 | 2062 | 1636 |
| 806 | 56497 | 49337 | 49221 | 49167 | 49116 | 573 |
| 808 | 30027 | 26125 | 26066 | 25985 | 25739 | 25336 |
| 809 | 39511 | 35067 | 35011 | 34986 | 34975 | 742 |
| 810 | 41395 | 36444 | 36361 | 36327 | 36327 | 2564 |
| 811 | 423 | 313 | 301 | 290 | 277 | 240 |
| 812 | 83865 | 76517 | 76358 | 74752 | 72215 | 72074 |
| 813 | 89338 | 82232 | 81950 | 81665 | 80557 | 80130 |
| 815 | 101310 | 92512 | 92334 | 90437 | 87601 | 87493 |
| 816 | 94189 | 85633 | 85446 | 84126 | 81946 | 81926 |
| 817 | 85367 | 77656 | 77492 | 77038 | 76095 | 76072 |
| 818 | 88556 | 80614 | 80415 | 79167 | 75474 | 75353 |
| 820 | 63887 | 58147 | 58079 | 56365 | 39229 | 39200 |
| 821 | 75692 | 68906 | 68727 | 68214 | 58129 | 57669 |
| 822 | 85184 | 74521 | 74336 | 73817 | 72091 | 63216 |
| 824 | 47400 | 42806 | 42714 | 41696 | 40909 | 32268 |
| 825 | 73851 | 67464 | 67398 | 67018 | 64837 | 44855 |
| 826 | 18053 | 15583 | 15541 | 15538 | 15538 | 1360 |
| 827 | 72227 | 64307 | 64097 | 63789 | 63583 | 41298 |
| 829 | 52892 | 47882 | 47805 | 47677 | 47093 | 30575 |
| 830 | 63085 | 56146 | 56069 | 56012 | 55929 | 41101 |
| 831 | 47012 | 42552 | 42467 | 42239 | 40448 | 23746 |
| 832 | 96545 | 88375 | 88274 | 87322 | 83532 | 83523 |
| 833 | 45186 | 41011 | 40878 | 39815 | 39653 | 39598 |
| 835 | 70635 | 64105 | 63746 | 63199 | 62236 | 62201 |
| 836 | 65216 | 59937 | 59656 | 59085 | 57390 | 57328 |
| 839 | 87529 | 79561 | 79327 | 78530 | 77753 | 77668 |
| 840 | 50566 | 45839 | 45764 | 45579 | 44934 | 44919 |
| 841 | 17591 | 16100 | 15982 | 15790 | 15130 | 15005 |
| 842 | 40010 | 35379 | 35294 | 35255 | 35250 | 6128 |
| 844 | 80261 | 72045 | 71895 | 71712 | 71017 | 42427 |
| 845 | 42271 | 37635 | 37532 | 37456 | 37376 | 26936 |
| 846 | 67476 | 60340 | 60273 | 60143 | 60022 | 30856 |
| 848 | 74606 | 66892 | 66742 | 66570 | 66442 | 38437 |
| 849 | 81751 | 70168 | 69659 | 69450 | 69434 | 8288 |
| 850 | 50874 | 44761 | 44565 | 44522 | 44522 | 115 |
| 851 | 28517 | 23701 | 23528 | 23499 | 23499 | 41 |
| 852 | 68115 | 61316 | 61206 | 61085 | 60781 | 60658 |
| 853 | 90420 | 82007 | 81864 | 81639 | 80980 | 80455 |
| 855 | 38075 | 34203 | 34099 | 33951 | 33417 | 33361 |
| 856 | 65321 | 58384 | 58282 | 57506 | 56565 | 56550 |
| 857 | 72890 | 65912 | 65766 | 65329 | 64257 | 64236 |
| 858 | 51516 | 45571 | 45491 | 45431 | 45431 | 385 |
| 859 | 63541 | 55630 | 55394 | 55288 | 55288 | 176 |
| 860 | 82790 | 73701 | 73517 | 73395 | 73171 | 30220 |
| 861 | 77666 | 69739 | 69550 | 69454 | 69360 | 51388 |
| 863 | 46801 | 41682 | 41576 | 41489 | 41489 | 12462 |
| 864 | 32634 | 28905 | 28833 | 28804 | 28804 | 90 |
| 865 | 52517 | 46854 | 46739 | 46662 | 44871 | 28093 |
| 866 | 26744 | 23971 | 23918 | 23900 | 23857 | 15549 |
| 867 | 80972 | 71598 | 71353 | 71271 | 71271 | 147 |
| 868 | 79462 | 72865 | 72777 | 72090 | 61682 | 61664 |
| 869 | 71792 | 66177 | 66040 | 65461 | 62969 | 62949 |
| 870 | 73832 | 67641 | 67451 | 67165 | 66591 | 66546 |
| 872 | 72781 | 66213 | 66082 | 65541 | 58299 | 58142 |
| 873 | 80260 | 73821 | 73609 | 73260 | 69459 | 69451 |
| 874 | 93324 | 85627 | 85380 | 84466 | 46378 | 46352 |
| 875 | 89635 | 80572 | 80373 | 78305 | 76283 | 76224 |
| 876 | 47759 | 43725 | 43633 | 43411 | 40150 | 39957 |
| 877 | 93029 | 84761 | 84683 | 83781 | 82462 | 82407 |
| 878 | 37996 | 33526 | 33436 | 33423 | 33306 | 4337 |
| 879 | 44393 | 39137 | 39078 | 39055 | 39053 | 469 |
| 880 | 44364 | 39105 | 39002 | 38970 | 38970 | 678 |
| 881 | 43940 | 39199 | 39111 | 39087 | 39058 | 11886 |
| 882 | 67005 | 58857 | 58781 | 58752 | 58752 | 60 |
| 884 | 46693 | 41333 | 41249 | 41231 | 41222 | 2976 |
| 885 | 84218 | 75557 | 75409 | 75378 | 75063 | 52926 |
| 886 | 16154 | 14552 | 14538 | 14525 | 14522 | 9081 |
| 888 | 18623 | 16350 | 16303 | 16265 | 16265 | 1310 |
| 890 | 93495 | 84536 | 84339 | 84020 | 83455 | 52304 |
| 891 | 33814 | 29154 | 29012 | 28968 | 28968 | 605 |
| 892 | 68899 | 61538 | 61353 | 61280 | 61269 | 36377 |
| 893 | 53019 | 47300 | 47083 | 47018 | 47006 | 7396 |
| 894 | 27940 | 24529 | 24393 | 24336 | 24336 | 2743 |
| 895 | 77142 | 68127 | 67942 | 67859 | 67764 | 18693 |
| 896 | 61401 | 54123 | 53947 | 53896 | 53896 | 191 |
| 897 | 36592 | 32251 | 32049 | 32003 | 32003 | 192 |
| 898 | 61140 | 56033 | 55809 | 55506 | 54715 | 54621 |
| 899 | 81010 | 73796 | 73524 | 73121 | 70937 | 70379 |
| 901 | 67213 | 61092 | 60927 | 60688 | 56697 | 56628 |
| 902 | 76521 | 68827 | 68418 | 68067 | 67220 | 48985 |
| 903 | 83012 | 76149 | 75876 | 75468 | 74658 | 74618 |
| 905 | 85591 | 78713 | 78334 | 77883 | 74108 | 74071 |
| 906 | 79791 | 72576 | 72400 | 72256 | 53633 | 53583 |
| 907 | 70041 | 64087 | 63822 | 63455 | 54228 | 54202 |
| 908 | 90885 | 82057 | 81927 | 81758 | 81417 | 55255 |
| 910 | 38514 | 33156 | 33105 | 33088 | 33088 | 147 |
| 911 | 55526 | 48612 | 48551 | 48522 | 48522 | 112 |
| 912 | 58413 | 51350 | 51191 | 51120 | 51120 | 504 |
| 914 | 32532 | 28688 | 28563 | 28525 | 28468 | 2789 |
| 915 | 50397 | 44553 | 44399 | 44337 | 44337 | 483 |
| 916 | 66433 | 58886 | 58674 | 58591 | 58571 | 3983 |
| 917 | 47203 | 41792 | 41729 | 41694 | 41694 | 173 |
| 918 | 78287 | 71256 | 71100 | 68976 | 66444 | 66394 |
| 919 | 30137 | 27358 | 27254 | 26882 | 26215 | 26198 |
| 920 | 88839 | 80685 | 80445 | 79952 | 78763 | 78662 |
| 922 | 13965 | 12560 | 12486 | 12134 | 11527 | 11496 |
| 923 | 78937 | 71728 | 71544 | 69733 | 65926 | 65901 |
| 924 | 50350 | 46064 | 45970 | 45720 | 45579 | 45502 |
| 925 | 81500 | 73348 | 73230 | 71114 | 58168 | 58132 |
| 926 | 59403 | 54162 | 54054 | 53211 | 48467 | 48461 |
| 927 | 9179 | 8276 | 8264 | 8258 | 8240 | 4548 |
| 928.1 | 12825 | 11327 | 11320 | 11316 | 11316 | 29 |
| 928.2 | 14290 | 12532 | 12526 | 12521 | 12521 | 93 |
| 929 | 94854 | 85061 | 84888 | 84822 | 84754 | 67974 |
| 930 | 3177 | 2761 | 2755 | 2740 | 2740 | 400 |
| 932 | 68820 | 62054 | 61964 | 61887 | 61788 | 24345 |
| 933 | 47999 | 42171 | 42068 | 42041 | 42003 | 1045 |
| 934 | 4055 | 3505 | 3485 | 3481 | 3481 | 210 |
| 936 | 75502 | 68632 | 68483 | 68345 | 67961 | 48645 |
| 937 | 39504 | 36175 | 35992 | 35811 | 35318 | 35067 |
| 938 | 75988 | 69335 | 69092 | 68574 | 61237 | 60994 |
| 940 | 98441 | 87336 | 87050 | 85946 | 84465 | 72637 |
| 941 | 72809 | 66729 | 66596 | 66043 | 59564 | 59531 |
| 943 | 74202 | 68253 | 67948 | 67619 | 65555 | 65542 |
| 945 | 74226 | 67670 | 67308 | 66689 | 65557 | 65312 |
| 946 | 59546 | 54633 | 54550 | 54396 | 53701 | 53684 |
| 947 | 30909 | 27398 | 27362 | 27349 | 27349 | 449 |
| 948 | 63077 | 55548 | 55416 | 55335 | 55108 | 30060 |
| 949 | 66075 | 60002 | 59936 | 59770 | 59736 | 41551 |
| 951 | 55140 | 48763 | 48658 | 48622 | 48622 | 266 |
| 952 | 279 | 213 | 192 | 183 | 183 | 0 |
| 953 | 44513 | 38669 | 38411 | 38298 | 38298 | 141 |
| 954 | 77552 | 68573 | 68336 | 68219 | 68219 | 7728 |
| 955 | 25011 | 20471 | 20415 | 20393 | 20381 | 7478 |
| 956 | 66516 | 58737 | 58578 | 58529 | 58529 | 148 |
| 957 | 33083 | 29078 | 29045 | 29020 | 29016 | 8096 |
| 959 | 78569 | 71612 | 71548 | 71492 | 71201 | 62019 |
| 960 | 84817 | 76398 | 76294 | 76188 | 62229 | 11214 |
| 961 | 66640 | 59281 | 59195 | 59162 | 59162 | 30 |
| 962 | 53605 | 47127 | 47068 | 47051 | 47051 | 122 |
| 963 | 241 | 148 | 126 | 107 | 107 | 0 |
| 964 | 53057 | 46409 | 46362 | 46345 | 46345 | 92 |
| 966 | 62235 | 54099 | 53946 | 53913 | 53913 | 110 |
| 967 | 58438 | 52740 | 52696 | 52641 | 50845 | 49081 |
| 968 | 82869 | 75386 | 75260 | 74633 | 65010 | 54303 |
| 969 | 68437 | 60620 | 60394 | 60289 | 60289 | 120 |
| 970 | 81792 | 72496 | 72345 | 72253 | 72247 | 316 |
| 971 | 80552 | 70815 | 70587 | 70470 | 70447 | 7061 |
| 973 | 19942 | 17460 | 17391 | 17380 | 17380 | 223 |
| 974 | 8866 | 6297 | 6264 | 6251 | 6251 | 23 |
| 975 | 57262 | 52161 | 52048 | 51902 | 51049 | 35175 |
| 976 | 107237 | 96393 | 96274 | 96208 | 96009 | 59314 |
| 977 | 67462 | 62151 | 62008 | 61537 | 58192 | 58179 |
| 978 | 75713 | 67991 | 67826 | 67459 | 65946 | 65946 |
| 979 | 97437 | 89128 | 89011 | 86342 | 84122 | 84104 |
| 981 | 87187 | 79859 | 79732 | 79319 | 73659 | 73657 |
| 982 | 18369 | 16805 | 16772 | 16720 | 16656 | 16646 |
| 983 | 76162 | 69383 | 69315 | 69155 | 67070 | 67055 |
| 984 | 56548 | 50956 | 50737 | 50093 | 48786 | 48765 |
| 985 | 5851 | 5085 | 5066 | 5043 | 5043 | 41 |
| 986 | 98876 | 90943 | 90845 | 88740 | 56582 | 56548 |
| NEG_1 | 508 | 58 | 32 | 18 | 18 | 0 |
| NEG_2 | 98 | 56 | 51 | 44 | 34 | 11 |
| NEG_3_1 | 130 | 9 | 1 | 0 | 0 | 0 |
| NEG_3_2 | 533 | 9 | 3 | 0 | 0 | 0 |

**Appendix A2 – Aggregation of data for estimation of** $\boldsymbol{A}_{\boldsymbol{t}}$

Our index of taxon/trait accessibility ($A_{t}$) is a ratio of proportions—proportion of a taxon/trait in the predator’s diet relative to the proportionate abundance of that taxon/trait in the environment. Before calculating $A_{t}$ we had to decide on the level of data aggregation that would comprise our individual observations (lowest resolution of data across which error is estimated; rows of the data frame). Individual fish guts had no obvious pairing with individual invertebrate samples. That is, fish were collected randomly throughout each sampling unit and are mobile, so likely predate across numerous areas within the sampling unit over the 48 h preceding capture (the period of predation arguably best represented by the gut contents at the time of capture (Chabot et al. 2016)). Further, to obtain $A_{t}$ for as many taxa/traits as possible (including rarer taxa) we had to aggregate a relatively large number of predator guts to estimate the proportionate abundance of a prey taxon/trait in the diet of each fish species/life-stage combination (following the recommendation of Kimmerer & Slaughter, 2021).

Therefore, our individual observations for estimates of $A_{t}$ were samples pooled across the three sampling units sampled within each month, within each year and reach. As such, an individual observation of $A_{t}$ involved pooling:

1. 39 invertebrate samples (3 sampling units x 13 samples per unit); and
2. up to 30 fish guts (3 sampling units x 10 fish per unit), noting that the 10 fish collected per sampling unit, per species, covered both life-stages (the total count of 30 was distributed as evenly as possibly across two life-stages).

**Appendix A3 – Estimating effects of environmental stressors on trait abundance**

Barrett et al. (2022) used non-metric multidimensional scaling (NMDS) to examine how the abundance of macroinvertebrate traits varied among rivers in Canterbury and the West Coast of New Zealand exposed to five stressors: flooding, drying, nutrient enrichment, fine deposited sediment, and acid mine drainage (AMD). Their NMDS produced three ordination axes that together described major gradients in *trait* abundance across sites. By correlating these axes with measured stressors, Barrett et al. (2022) identified which stressor gradients each ordination axis represented, and therefore which stressor conditions each *trait* would be most abundant in. For example, NMDS axis 1 was positively correlated with deposited fine sediment, and the ≤5 mm body-size *trait* had the highest NMDS scores along this axis relative to other body size *traits*—indicating that small-bodied taxa were most abundant under conditions of high sediment stress.

This information was presented graphically by Barrett et al. (2022). We therefore developed an algorithm to translate this information into a quantitative measure of the relative abundance of each *trait* under stressful conditions of each stressor. The procedure involved four main steps for each stressor:

1. We identified all NMDS axes that were significantly correlated to the stressor and noted the direction of the relationship reported by Barrett et al. (2022). For example, all three NMDS axes were positively related to AMD, meaning that higher axis scores corresponded to more stressful AMD conditions.
2. For each of these NMDS axes, we ranked *trait*s within each *trait group* according to their NMDS axis scores. *Traits* with NMDS axis scores that were associated with the most stressful conditions of a stressor were given the highest numerical rank. For instance, the ≤5 mm body-size *trait* had the highest scores for NMDS axis 1 among all *traits* within the maximum body size *trait group*. Because NMDS axis 1 was positively correlated with AMD stress, this implied that ≤5 mm sized taxa were the most abundant in AMD-stressed sites, relative to other sized taxa. Consequently, the ≤5 mm body-size *trait*, received rank 5, while the remaining four *traits* within the body size trait group received lower ranks. Trait NMDS axis scores were visually extracted from Figures 3-5 in Barrett et al. (2022).
3. We then calculated the median rank of each *trait* across all NMDS axes that were significantly related to the stressor. For example, the ≤5 mm size *trait* received ranks 5, 5, and 2 for NMDS axes 1, 2 and 3, respectively, giving a median rank of 5.
4. Finally, we normalised the median rank by dividing it by the maximum rank within a *trait group*, producing a value between 0 and 1. The resulting value, *T_iks_*, was the relative abundance of *trait* *i*, in rivers impacted by stressor *s*, among *traits* in *trait group* *k.* Values of 1 indicated the *trait* was highly abundant under stressful conditions of a stressor (i.e., high deposited fine sediment cover, nutrient enrichment and AMD and larger and more frequent drying and flooding events).

The stressor gradients used by Barrett et al. (2022) were as follows. Flooding intensity was measured using the Pfankuch River Disturbance Index (Pfankuch 1975; McHugh et al. 2010), where higher values indicate more frequent or severe high-flow disturbance, hence worsening flooding stress. Drying intensity was quantified as wetted cross-sectional channel area, with smaller areas reflecting greater drying stress (McHugh et al. 2015). Eutrophication was defined by a principle components analysis (PCA) axis combining nitrate and phosphorous concentrations, gross primary productivity, macrophyte cover, and shade (Graham et al. 2015), with higher scores indicating greater eutrophication stress. Fine deposited sediment was estimated visually as percent cover of particles <2 mm with higher cover considered more stressful (Burdon et al. 2013). Finally, the AMD gradient was described by a PCA axis derived from pH, conductivity, and dissolved metal concentrations (Pomeranz et al. 2019), with higher scores reflecting greater stress. Associated macroinvertebrate community samples were taken using a Surber sampler in all studies.

|  | Body Form | | | | Body Flexibility | | | Maximum Size | | | | | Feeding Habit | | | | | | Substrate Attachment | | | |
| --- | --- | --- | --- | --- | --- | --- | --- | --- | --- | --- | --- | --- | --- | --- | --- | --- | --- | --- | --- | --- | --- | --- |
| Taxon | Cylindrical | Flattened | Spherical | Streamlined | None | Moderate | High | ≤5 mm | >5–10 mm | >10–20 mm | >20–40 mm | >40 mm | Algal | Deposit | Filter | Scraper | Shredder | Predator | Attached | Burrower | Crawler | Swimmer |
| Archichauliodes | 0.5 | 0.5 | 0 | 0 | 0 | 0.75 | 0.25 | 0 | 0 | 0 | 0 | 1 | 0 | 0 | 0 | 0 | 0 | 1 | 0 | 0 | 1 | 0 |
| Atalophlebioides | 0 | 0.75 | 0 | 0.25 | 0 | 1 | 0 | 0 | 1 | 0 | 0 | 0 | 0 | 0 | 0 | 1 | 0 | 0 | 0 | 0 | 1 | 0 |
| Chironomidae | 1 | 0 | 0 | 0 | 0 | 0 | 1 | 0.33 | 0.33 | 0.33 | 0 | 0 | 0 | 0.14 | 0 | 0.45 | 0 | 0.41 | 0 | 0.5 | 0.47 | 0.03 |
| Clitellata | 0.89 | 0.11 | 0 | 0 | 0 | 0 | 1 | 0 | 0.11 | 0.11 | 0.33 | 0.44 | 0 | 0.86 | 0 | 0 | 0 | 0.14 | 0 | 0.81 | 0.1 | 0.1 |
| Coloburiscus | 1 | 0 | 0 | 0 | 0 | 1 | 0 | 0 | 0 | 1 | 0 | 0 | 0 | 0 | 1 | 0 | 0 | 0 | 0.6 | 0 | 0.4 | 0 |
| Deleatidium | 0 | 0.75 | 0 | 0.25 | 0 | 1 | 0 | 0 | 0.62 | 0.38 | 0 | 0 | 0 | 0 | 0 | 1 | 0 | 0 | 0 | 0 | 1 | 0 |
| Hydrobiosis | 0.86 | 0.07 | 0 | 0.07 | 0 | 0.6 | 0.4 | 0 | 0 | 0.62 | 0.38 | 0 | 0 | 0 | 0 | 0 | 0 | 1 | 0 | 0 | 1 | 0 |
| Hydropsychidae | 1 | 0 | 0 | 0 | 0 | 0.75 | 0.25 | 0 | 0.18 | 0.82 | 0 | 0 | 0 | 0 | 0.75 | 0 | 0 | 0.25 | 0.67 | 0 | 0.33 | 0 |
| Muscidae | 1 | 0 | 0 | 0 | 0 | 0 | 1 | 0 | 0 | 1 | 0 | 0 | 0 | 0 | 0 | 0 | 0 | 1 | 0 | 0 | 1 | 0 |
| Neurochorema | 1 | 0 | 0 | 0 | 0 | 0.6 | 0.4 | 0 | 0.25 | 0.75 | 0 | 0 | 0 | 0 | 0 | 0 | 0 | 1 | 0 | 0 | 1 | 0 |
| Olinga | 1 | 0 | 0 | 0 | 1 | 0 | 0 | 0 | 1 | 0 | 0 | 0 | 0 | 0 | 0 | 0.75 | 0.25 | 0 | 0 | 0 | 1 | 0 |
| Ostracoda | 0 | 0.75 | 0.25 | 0 | 1 | 0 | 0 | 1 | 0 | 0 | 0 | 0 | 0 | 0.33 | 0 | 0.22 | 0.22 | 0.22 | 0 | 0.17 | 0.5 | 0.33 |
| Oxyethira | 0.4 | 0.6 | 0 | 0 | 0.25 | 0.75 | 0 | 1 | 0 | 0 | 0 | 0 | 0.56 | 0 | 0 | 0.44 | 0 | 0 | 0 | 0 | 1 | 0 |
| Physella | 1 | 0 | 0 | 0 | 1 | 0 | 0 | 0 | 0 | 1 | 0 | 0 | 0 | 0 | 0 | 1 | 0 | 0 | 0 | 0 | 1 | 0 |
| Potamopyrgus | 1 | 0 | 0 | 0 | 1 | 0 | 0 | 0 | 0 | 1 | 0 | 0 | 0 | 0 | 0 | 1 | 0 | 0 | 0 | 0 | 1 | 0 |
| Psilochorema | 0.76 | 0 | 0 | 0.24 | 0 | 0.6 | 0.4 | 0 | 0 | 1 | 0 | 0 | 0 | 0 | 0 | 0 | 0 | 1 | 0 | 0 | 1 | 0 |
| Pycnocentria | 1 | 0 | 0 | 0 | 1 | 0 | 0 | 0 | 1 | 0 | 0 | 0 | 0 | 0 | 0 | 0.44 | 0.56 | 0 | 0 | 0.2 | 0.8 | 0 |
| Pycnocentrodes | 1 | 0 | 0 | 0 | 1 | 0 | 0 | 0 | 1 | 0 | 0 | 0 | 0 | 0 | 0 | 1 | 0 | 0 | 0 | 0 | 1 | 0 |
| Simuliidae | 1 | 0 | 0 | 0 | 0 | 0 | 1 | 0 | 1 | 0 | 0 | 0 | 0 | 0 | 0.75 | 0.25 | 0 | 0 | 1 | 0 | 0 | 0 |

**Table S1:** Standardised affinity scores $\left( \tilde{a}_{ikj} \right)$for prey taxa used in to calculate trait abundance in gut and benthic samples.

**Table S2**: Sources and credits for silhouettes used from Phylopic to represent some taxa in figures.

| Taxon | Phylopic URL | Credit |
| --- | --- | --- |
| Chironomidae | [Chironomidae by Nico Muñoz (CC BY-NC 3.0) - PhyloPic](https://www.phylopic.org/images/834f9ef5-c5bf-4e9e-94c8-3ecb8fb14838/chironomidae) | Nico Muñoz |
| Clitellata | [Lumbricus terrestris (CC0 1.0) - PhyloPic](https://www.phylopic.org/images/f0d839a1-5a3e-42b2-828a-c5e536f0f7b7/lumbricus-terrestris) | Carlo De Rito |
| Ostracoda | [Cypris by Maxime Dahirel (CC BY 3.0) - PhyloPic](https://www.phylopic.org/images/8d3b12ff-02f6-4b7d-8387-2b3374f9d393/cypris) | Maxime Dahirel |
| Simuliidae | [Simuliidae by Nico Muñoz (CC BY-NC 3.0) - PhyloPic](https://www.phylopic.org/images/b6264258-cee4-4cd8-85ef-ea03d127204f/simuliidae) | Nico Muñoz |
| Physella | [Physella acuta by S. Ginot (CC0 1.0) - PhyloPic](https://www.phylopic.org/images/6cb3954e-03cc-4121-85e8-1a083b70706c/physella-acuta) | Samuel Ginot |
